# Structural Insights into the DNA-Binding Mechanism of BCL11A: The Integral Role of ZnF6

**DOI:** 10.1101/2024.01.17.576058

**Authors:** Thibault Viennet, Maolu Yin, Abhilash Jayaraj, Woojin Kim, Zhen-Yu J. Sun, Yuko Fujiwara, Kevin Zhang, Davide Seruggia, Hyuk-Soo Seo, Sirano Dhe-Paganon, Stuart H. Orkin, Haribabu Arthanari

**Affiliations:** Department of Cancer Biology, Dana-Farber Cancer Institute, Boston, MA, USA; Biological Chemistry and Molecular Pharmacology, Harvard Medical School, Boston, MA, USA; Dana Farber/Boston Children’s Hospital Cancer and Blood Disorders Center, Boston, MA, USA; Howard Hughes Medical Institute, Boston, MA, USA; Department of Pediatrics, Boston Children’s Hospital, Harvard Medical School, Boston, MA, USA; Department of Chemistry and iNANO, Aarhus University, Aarhus, Denmark

## Abstract

The transcription factor BCL11A is a critical regulator of the switch from fetal hemoglobin (HbF: α_2_γ_2_) to adult hemoglobin (HbA: α_2_β_2_) during development. BCL11A binds at a cognate recognition site (TGACCA) in the γ-globin gene promoter and represses its expression. DNA-binding is mediated by a triple zinc finger domain, designated ZnF456. Here, we report comprehensive investigation of ZnF456, leveraging X-ray crystallography and NMR to determine the structures in both the presence and absence of DNA. We delve into the dynamics and mode of interaction with DNA. Moreover, we discovered that the last zinc finger of BCL11A (ZnF6) plays a special role in DNA binding and γ-globin gene repression. Our findings help account for some rare γ-globin gene promoter mutations that perturb BCL11A binding and lead to increased HbF in adults (hereditary persistence of fetal hemoglobin). Comprehending the DNA binding mechanism of BCL11A opens avenues for the strategic, structure-based design of novel therapeutics targeting sickle cell disease and β-thalassemia.

## Introduction

Inherited disorders of hemoglobin, Sickle Cell Disease (SCD) and β-thalassemia, are among the most prevalent genetic diseases in the world with greater than 300,000 babies born each year, the majority in resource-limited countries^1^. These disorders result from mutations in the adult β-globin gene. In SCD, a single amino acid substitution alters the properties of β-globin and promotes polymerization (sickling) of the hemoglobin tetramer (α_2_β^S^_2_). In β-thalassemia, mutations impair β-globin production, leading to deficiency of adult hemoglobin (HbA: α_2_β_2_)^2^. Early on, it was recognized that elevated levels of fetal hemoglobin (HbF, α_2_γ_2_) counteracts severity of both conditions^3,4^. This recognition prompted investigations into mechanisms by which expression of γ-globin during fetal life is silenced and replaced by β-globin^5^. Genome-wide association studies led to discovery of BCL11A as a potential γ-globin silencer^6,7^. Subsequent studies have validated this role, defined its direct action at the γ-globin gene, and led to successful clinical trials in both SCD and β3-thalassemia^8-11^.

Three C-terminal zinc finger domains (ZnF456) of BCL11A mediate its sequence specific binding to DNA^12,13^ to an optimal site TGACCA^14,15^. This sequence is duplicated in the promoters of the γ-globin genes. Indeed, BCL11A binds specifically to the distal duplicate, which is located at -118 to -113. Remarkably, rare single base substitutions within this region occur in individuals with hereditary persistent expression of HbF (HFPH syndrome) as adults^13,16^. These mutations impair binding of BCL11A to the promoter^14,17^.

A 2.5 Å resolution crystal structure of the C-terminal three zinc fingers of BCL11A (aa 731-835) bound to a 12-bp double-stranded DNA oligo of sequence CCTTGACCAATA, corresponding to the -121 to -110 γ-globin promoter region, has been reported^17^. It was observed that the first two zinc fingers (ZnF45) bind to the major groove of DNA, while the last zinc finger (ZnF6) had poor electron density and rare contacts with DNA, well in line with a lower contribution of ZnF6 to DNA binding seen previously in cells^13^.

Here, we extend these findings and report on the dynamics and structure of BCL11A in the apo form and bound to a 19-bp HPHF γ-globin promoter region using a combination of X-ray crystallography and nuclear magnetic resonance (NMR). Additionally, we characterize the special role of zinc finger 6 in DNA binding and γ-globin gene repression.

## Results

### Crystal structure of BCL11A bound to a 19-bp DNA

Here we report the X-ray crystallographic structure of BCL11A ZnF456 in complex with DNA. We initially determined the structure of ZnF456 bound to a 12-bp double-stranded oligonucleotide (PDB code 6U9Q) and the protein structure is virtually identical to a prior study of Yang *et al*.^17^. In their report, the last zinc finger (ZnF6) bound in the minor groove of DNA with no direct contacts, but its electron density was not as well defined with ZnF6 being missing in one of the two complexes in the crystal unit. We hypothesized that this could be a result of the shorter DNA used, potentially preventing ZnF6 from adopting its natural bound state. We therefore determined the structure of ZnF456 bound to the longer DNA sequence.

We obtained the X-ray crystal structure of BCL11A ZnF456 (aa 730-835) bound to a 19-bp DNA oligonucleotide at 1.8 Å resolution, including for ZnF6. Coordinates were deposited in the PDB under accession number 8TLO (see Table S1 for structure statistics). The zinc fingers have simple, modular structures, with each zinc finger domain comprised of an antiparallel β-sheet and an α-helix, which are held together by a zinc ion and a series of hydrophobic residues. The three zinc fingers are arranged in a semicircular structure that fits tightly into the DNA major and minor grooves. Zinc finger 4 (ZnF4) and zinc finger 5 (ZnF5) bind to the major groove, while zinc finger 6 (ZnF6) engages the minor groove as depicted in Fig. 1A. It is notable that the α-helices are tipped at a steeper angle than the angles of the grooves.

**Figure 1.**
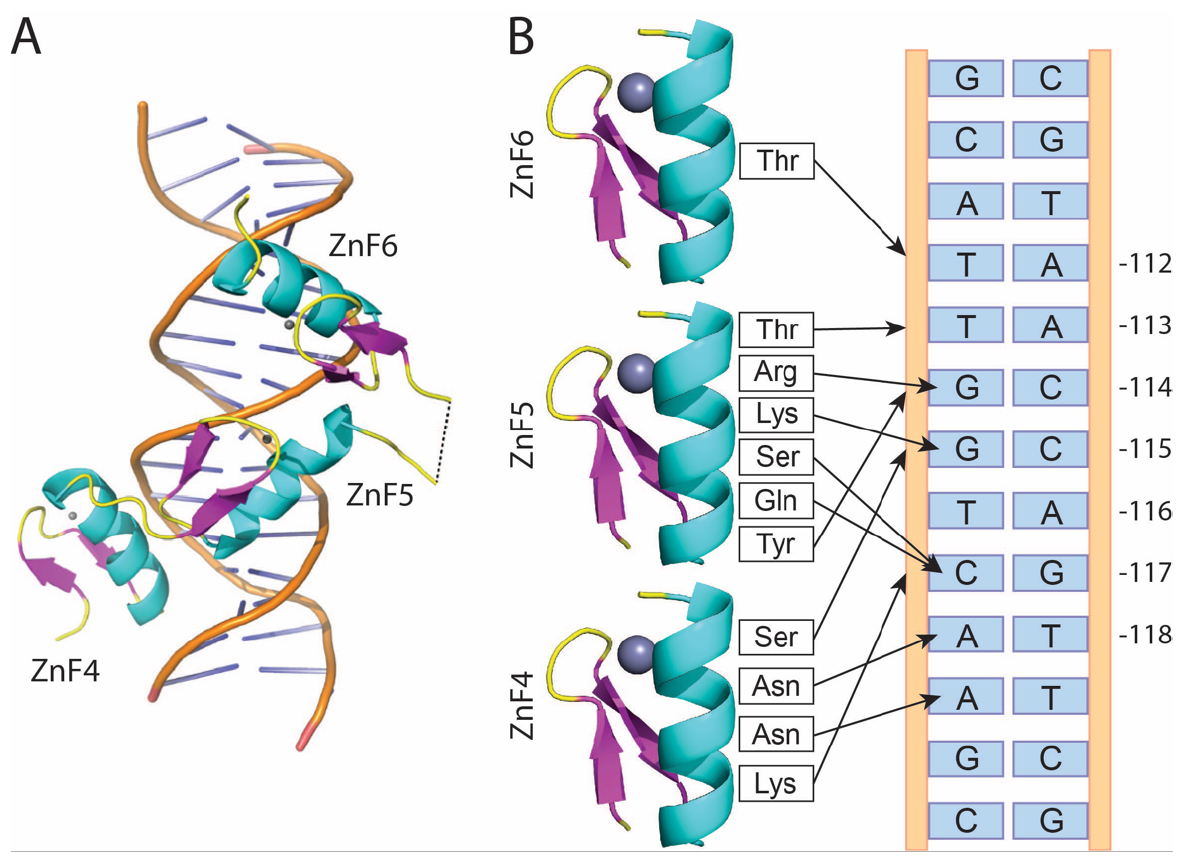
X-ray structure of BCL11A bound to DNA reveals the role of each zinc finger domain. A) 1.8 Å X-ray crystal structure of ZnF456 bound to a 19-bp DNA oligo. Protein is shown as ribbon cartoon. Helices are colored in cyan, β-sheets in magenta and coils in yellow. B) Representation of the residues involved in specific (arrow to the DNA base) and unspecific (arrow to the DNA backbone) contacts of ZnF456 with DNA.

In greater detail, the BCL11A protein-DNA complex structure reveals a characteristic pattern of contacts of the protein α-helices with either the nucleotide, the DNA backbone, or a combination of both. The ZnF456 motif makes eleven critical contacts with the DNA helix that are described in Fig. 1B. ZnF4 has four contacts with the DNA, with two one of them contributing to sequence specificity. Asn753 forms a hydrogen bond with the adenine nucleotide at position - 119 and Asn756 with the adenosine at -118. Lys749 and Ser763 bind to the DNA backbone. While these do not contribute to specificity, they strengthen the protein-DNA interaction and place ZnF4 as a major driver of DNA binding. However, ZnF5 is the most critical part of the protein in the binding process with six contacts to DNA, including four specific contacts with nucleotides. Gln781 and Ser783 both contact the cytosine nucleotide at position -117, Lys784 and Arg787 bind to the guanine at -115 and -114, respectively, which contribute heavily to the sequence specificity of BCL11A to this DNA sequence. Additionally, Tyr777 and Thr791 bind to the DNA backbone and contribute to affinity. ZnF6 has no residues that contribute to sequence specificity (no direct contacts to the nucleobases), but Thr814 also binds to the DNA backbone. The β-sheets are on the back of the helices away from the base pairs, but it is notable that ZnF4 and ZnF5 β-sheets contribute to binding affinity through Lys749 and Tyr777, respectively.

The complex structure of ZnF456 bound to the 19-bp DNA reported here is very close to that of ZnF456 bound to the 12-bp DNA that was also determined by Yang *et al*.^17^, confirming that ZnF6 indeed naturally binds DNA in the minor groove and does not contribute to sequence specificity. Although it was suggested that it does not contribute to the affinity of BCL11A for the γ-globin promoter region^17^, there is one contact to the DNA backbone participating in the interaction.

#### Solution structure of apo BCL11A reveals specific conformational plasticity

While many DNA-bound tandem zinc finger repeat structures have been determined and exhibit similar arrangements^18-20^, much less is known about their apo structures. Knowledge of DNA-free apo structures may prove valuable for drug discovery efforts, which have been extremely challenging in targeting transcription factors^21^. Due to the high intrinsic plasticity of apo BCL11A ZnF456 and its failure to form crystals, we used NMR to determine the apo structure in solution.

Using conventional NMR approaches, we assigned 95% of non-proline backbone resonances and 82% of all sidechain resonances (BMRB ID: 52029). All secondary structure elements have high structural order parameters S^2^ above 0.8, as predicted by chemical shifts (Fig. 2A-B). This establishes that BCL11A C-terminal zinc fingers are well folded even in the absence of DNA. Interestingly, the linkers connecting ZnF4/5 and ZnF5/6 have different behaviors in terms of order parameters, with S^2^ in the range of 0.5 and 0.3, respectively. This is in line with the fact that while the linker connecting ZnF4/5 has the canonical TGEK/RP sequence^22^, the linker connecting ZnF5/6 has an unusual GQVGKDV that is longer, Gly-rich and devoid of Proline residues, and hence more flexible.

**Figure 2.**
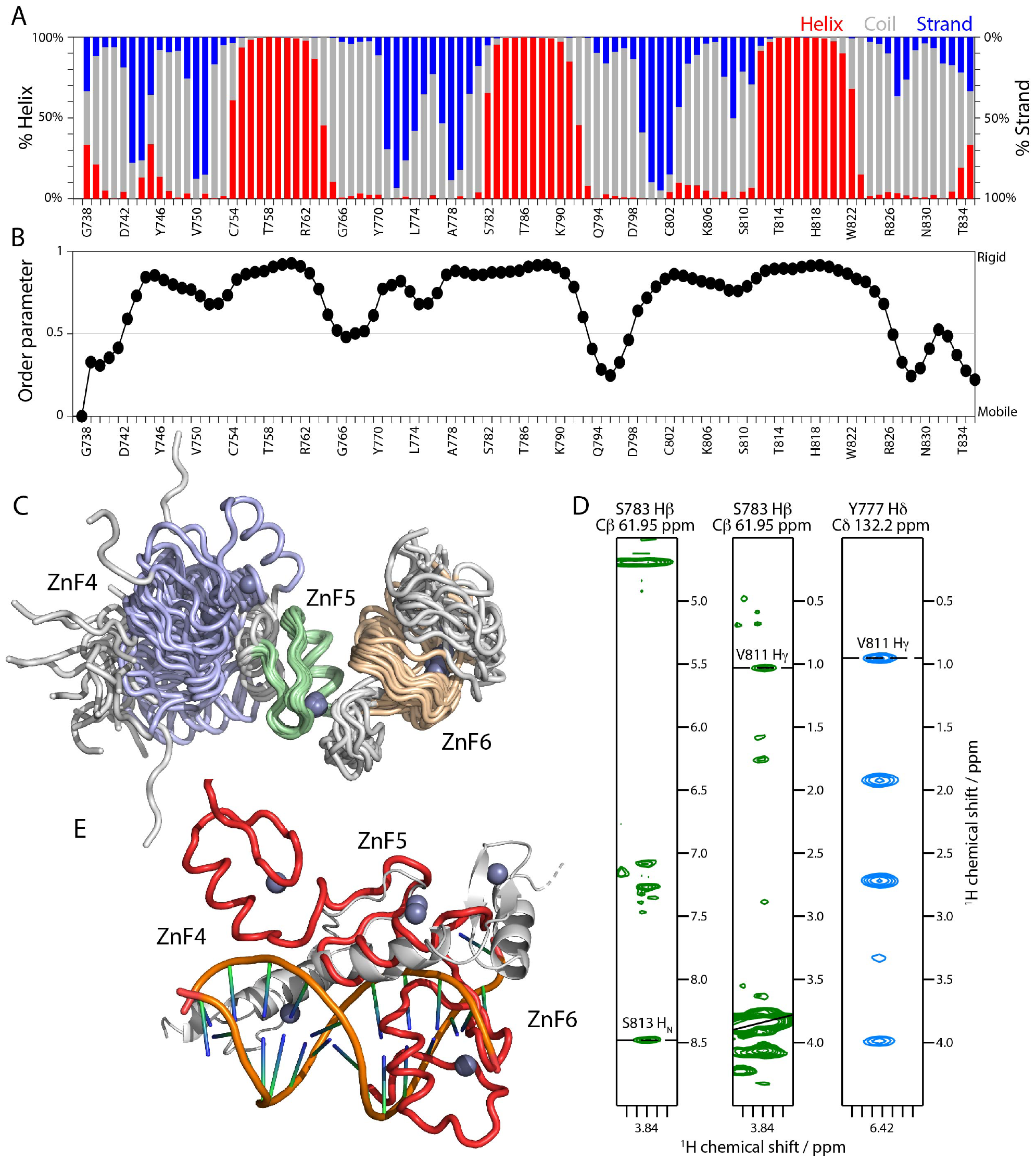
NMR solution structure of apo ZnF456 reveals specific contacts between ZnF5/6. A) Secondary structure propensity prediction based on chemical shifts for each residue of ZnF456 by TALOS-N. Red represents α-helices, blue represents β-strands and grey represents random coil or turn. B) Corresponding chemical shift-based residue-specific order parameters. C) NMR solution structural ensemble (20 lowest energy conformers after energy minimization) aligned on ZnF5. D) Select 3D NOE strip plots showing distance restraints between ZnF5 and ZnF6. E) Overlay of DNA-bound crystal structure (grey) and average solution structure (red), aligned on ZnF5.

We combined dihedral angles derived from chemical shifts, NOE cross-peaks, and hydrogen bond restraints, with H-N residual dipolar couplings (RDCs) to determine the solution “structure” (conformational ensemble) of DNA-free ZnF456 in the presence of Zn^2+^ (PDB ID: 8THO). The solution structure of each individual ZnF is well defined with backbone rmsd ranging from 0.39 to 0.81 Å, however the full-length ZnF456 has a backbone rmsd of 4.06 Å, in line with a high degree of conformational plasticity (Fig. 2C, detailed structural statistics in Table S2).

Notably, while ZnF4 and ZnF5 do not appear to be correlated in the conformational ensemble, ZnF5 and ZnF6 adopt a set of preferential orientations relative to one another. This is supported by the presence of a set of long-range NOEs between residues 777-783 in ZnF5 and residues 811-813 in ZnF6 (Fig. 2D). However, because NOEs are insensitive in dynamic conformational ensembles and prone to assignment errors, we analyzed our RDC data to quantify the degree of correlation between zinc fingers. RDCs report on the orientation of chemical bonds in respect to an alignment tensor and therefore are sensitive probes for the orientation of domains to one another. To determine the structural ensemble, we used a different tensor for each zinc finger which resulted in a very good fit quality (Q < 20%). The fit quality of RDC data to tensors calculated for pairs of two fingers report on the degree of correlation between them. Indeed, the fit quality is significantly better for ZnF5/6 (Q = 56%) compared to ZnF4/5 (Q = 63%) supporting the notion that ZnF5 and 6 have long range interactions as represented by specific orientational restraint between zinc fingers 5 and 6 (Table S3). This could be counterintuitive given the higher flexibility of the linker connecting ZnF5/6 but the higher degree of freedom allows specific domain-domain interactions, hence the correlation of ZnF5 and 6 is dictated by domain structures and interactions rather than by linker sequence and flexibility^22^.

We performed classical unbiased molecular dynamics simulation of ZnF456 starting from the X-ray structure but in the absence of DNA. The simulation protocol including the force field was modified to account for the intrinsically disordered nature of ZnF456 and is described in detail in the methods section. While the MD simulation demonstrated a high degree of conformational plasticity, simulation of ZnF456 in absence of DNA showed that ZnF5 is more corelated to ZnF6 than ZnF4 (Fig. S1A). This matches with our NMR results (Fig. 2C). Additionally, we assessed the variability in dihedral angles between our MD simulation trajectory and NMR ensemble (Fig. S1B,C). The linker residues 765-769 (ZnF4-ZnF5) and 793-799 (ZnF5-ZnF6) impart BCL11A its conformational flexibility in both MD and NMR structures (Fig. S1B,C).

The comparative analysis of the DNA-free (solution) and DNA-bound (crystal) states of BCL11A reveals a significant conformational rearrangement in the relative positioning of zinc fingers during DNA engagement. This conformational shift is visually presented in a morphing movie provided in the Supplementary Information. In solution, BCL11A adopts a structure that is incompatible to bind DNA, primarily due to spatial conflicts between ZnF6 and DNA if ZnF5 is used as an alignment reference, as depicted in Figure 2E. NMR data of the unbound BCL11A indicate that within the apo structural ensemble, ZnF5 and ZnF6 exhibit limited flexibility in respect to one another. This positions ZnF6 as a crucial element for DNA interaction, notwithstanding its smaller contribution to DNA binding affinity^13^. This nuanced understanding of ZnF6’s role in DNA recognition could provide insights into the molecular basis of BCL11A’s selective binding.

#### BCL11A dynamics informs its DNA binding mode

To elucidate the binding mode and dynamics of BCL11A ZnF456 to DNA we used NMR based relaxation experiments which provide a more detailed view of the dynamics that informs on structural rearrangements that occur upon binding.

We measured R_1_ and R_2_ relaxation rates as well as heteronuclear Overhauser effect (hetNOE) on apo ZnF456. The product of R_2_ and R_1_ (R_2_·R_1)_ relaxation rates serves as an indicator of the S^2^ dynamic order parameter^23^. Only the termini and the linker connecting ZnF5/6 have sizably lower S^2^ than the rest of the protein (Fig. 3A and Fig. 1A-B, black). The same trend is visible in the hetNOE, with zinc finger domains having hetNOE between 0.6 and 1 (rigid) and linkers having hetNOE values below 0.6, as low as 0.2 for the loop connecting ZnF5/6 (Fig. S2C, black). This confirms that the degree of freedom between ZnF5/6 is higher than between ZnF4/5, despite the specific orientations of ZnF5 and ZnF6 in respect to each other, clearly indicating that the apo structural ensemble is dictated by domain-domain interactions rather than by linker sequence and flexibility^22^.

**Figure 3.**
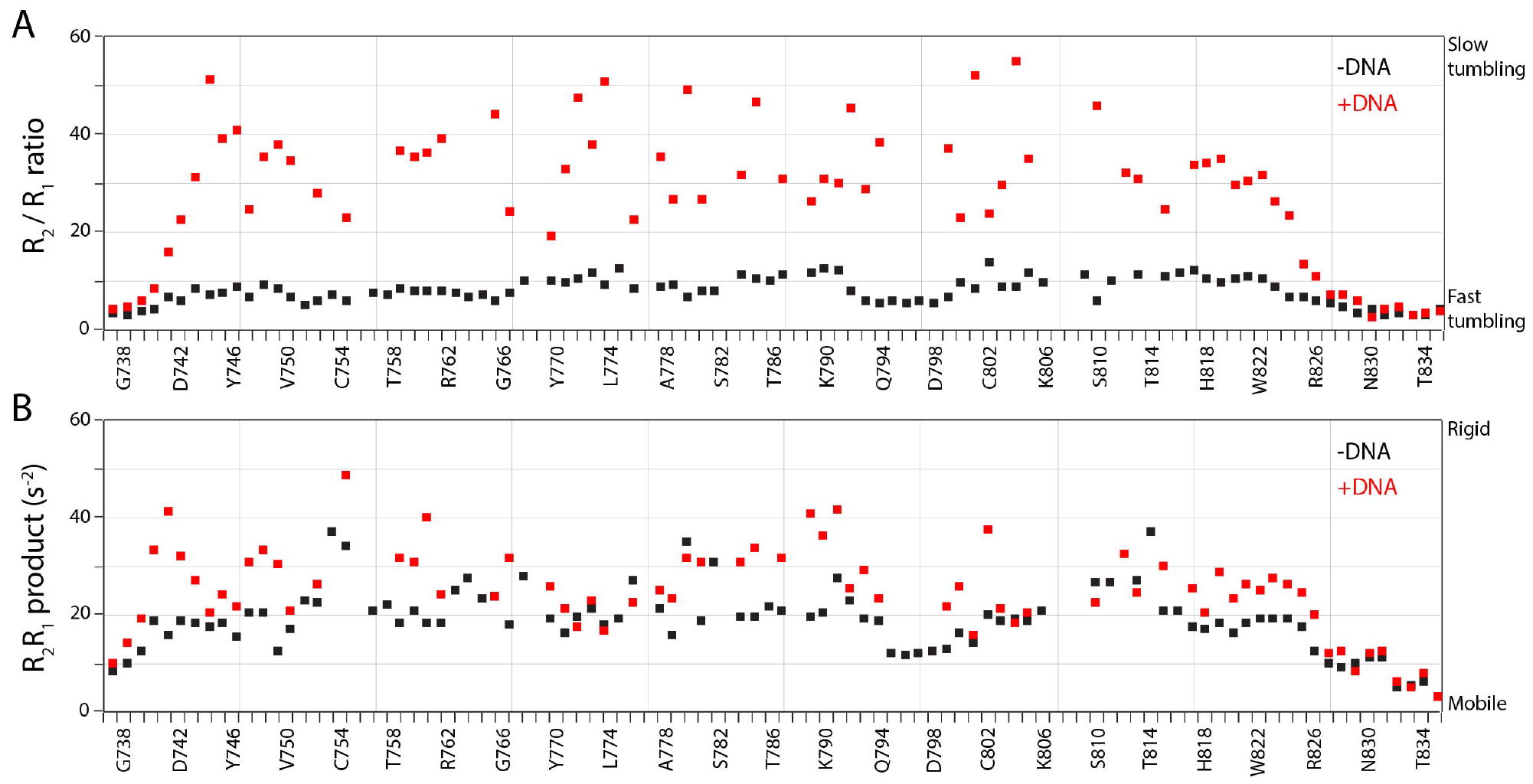
NMR sheds light on the dynamic process of BCL11A binding to DNA. A) Ratio of transverse (R_2_) and longitudinal (R_1_) relaxation rates for each residue of ZnF456 in the absence (black) and presence (red) of DNA, measuring tumbling time in solution. B) Product of transverse (R_2_) and longitudinal (R_1_) relaxation rates for each residue of ZnF456 in the absence (black) and presence (red) of DNA, proportional to the dynamic order parameter.

To address the seemingly paradoxical nature of these observations, we further investigated the dynamics using a truncated version of the protein lacking ZnF6. We observed no substantial chemical shift perturbations in the ZnF45 construct compared to the ZnF456, except for the newly exposed C-terminal residues (refer to Figure S3A-B). However, the R_2_ relaxation rates for ZnF5 were notably reduced (as seen in Figure S3C), which supports the notion of increased ZnF5 mobility in the absence of stabilizing contact with ZnF6. These insights are consistent with the hypothesis that ZnF5 exhibits greater mobility when not in contact with ZnF6, contributing to our understanding of the protein’s dynamic behavior in DNA binding.

We then complexed ZnF456 with a 14-bp double stranded DNA fragment corresponding to the distal γ-globin promoter region^13^. The NMR data (^1^H-^15^N-HSQC spectrum) showed extensive chemical shift perturbations and expected line broadening consistent with the formation of a larger complex (Fig. S4). Notably, these spectral shifts were pronounced in specific regions: the first β-strand and α-helix of ZnF4, the second β-strand and α-helix of ZnF5, and the second β-strand and initial segment of the α-helix in ZnF6. These observations suggest that ZnF6 also contributes binding to DNA, in contrast with the prior hypothesis^17^. It is important to recognize that the observed chemical shift perturbations not only capture direct interactions with DNA, but also the intricate conformational adaptations of the protein upon complexation with DNA.

Comparing the relaxation rates of apo ZnF456 and in the complex provides more detailed insights. The ratio R_2_/R_1_ is proportional to the effective tumbling time τ_C_ in solution^23^ and should correlate with the size of the complex in case of tight binding. While in the absence of DNA, ZnF456 has ratios around 5 (corresponding to a τ_C_ of 5.5 ns, and a size of approx. 9 kDa), in the presence of DNA it is elevated to an average of 25 (corresponding to a τ_C_ of 13.6 ns, and a size of approx. 23 kDa) (Fig. 3B and Fig. S2A-B). This finding aligns with a tight binding mode and the size of the complex. Only the N and C terminal residues do not have elevated R_2_/R_1_ ratios and there are no significant differences between other residues, demonstrating that all zinc fingers are tightly bound to DNA and tumble in solution at the size of the complex.

Dynamic order parameters (approximated by R_2_·R_1_) show a more nuanced behavior (Fig. 3A and Fig. S2A-B). Termini and linkers retain similar S^2^ as in the absence of DNA, whereas structured regions tend to have elevated order parameters, proving that they rigidify in the presence of DNA. It is particularly true for ZnF5 (+10 s^-2^ in R_2_·R_1_) and less prominent for ZnF6 (+6 s^-2^). Heteronuclear NOEs do not show any significant changes apart from a slight rigidification of ZnF5 (Fig. S2C). These observations have two major implications: (i) there may be a high entropic cost associated with binding ZnF5 to DNA but less for ZnF6 and (ii) the linker connecting ZnF5/6 retains its flexibility in the complex, potentially allowing the extensive conformational rearrangements required to avoid ZnF6 clashing with DNA.

We next conducted CPMG relaxation dispersion experiments that reports on slower motions (μs to low-ms time scale) on both the free and DNA-bound ZnF456 (Fig. S5). Statistical Monte Carlo analysis of dispersion curves did not detect any motions on the μs-ms timescale except for specific residues in the free BCL11A: Glu803, Thr814, and Arg826, all of which are part of ZnF6. This could be attributed to a conformational exchange occurring between the states when ZnF5 is bound and when it is not. However, the analysis did not detect any correlated movements typically indicative of a conformational selection mechanism, where a pre-existing conformation is selected from a repertoire of states upon ligand binding.

To further our understanding of the binding process of BCL11A to DNA, we used MD to simulate the binding process. No restraints were employed to artificially direct BCL11A towards the DNA. Remarkably, one of the starting configurations spontaneously formed a stable complex with the DNA that persisted for over 2 microseconds. Analysis of the trajectory revealed that ZnF5 is the first to interact with DNA followed by ZnF6 and subsequently by ZnF4. This is in agreement with previous observations that ZnF5 is critical to DNA binding process and point mutations in ZnF5 helix leads to loss in binding affinity^17^. Additionally, we observed transient hydrogen bonds between ZnF6 and DNA in the early stages of binding and recognition phase. Positively charged residues Lys817, Lys820, Lys821 of ZnF6 contact negatively charged backbone nucleotides of minor groove DNA. These contacts assist ZnF5 to achieve its correct alignment within the major groove of DNA. ZnF4, however, was flexible even after initial binding of BCL11A and was later stabilized during the simulation by sequence specific nucleotide contacts.

#### A specific role for zinc finger 6 in DNA binding

In light of the DNA-bound crystal structure, the DNA-free solution structure and our dynamics data upon DNA binding, we conclude that ZnF6 plays a special role in DNA recognition. In the solution structure, ZnF6 is in a conformation that would clash with DNA upon binding, thus it reorients itself in respect to ZnF5 to allow binding. This is allowed by its higher degree of freedom in the solution conformational ensemble due to the non-canonical linker between ZnF5 and ZnF6. We explored the consequences of this special role of ZnF6 for BCL11A.

We measured the affinity of ZnF456 (*K*_d_ = 19 ± 4 nM) and ZnF45 (lacking zinc finger 6, *K*_d_ = 225 ± 19 nM) to DNA using isothermal titration calorimetry (Fig. 4A). ZnF6 presence increases the affinity by 12-fold, therefore contributing the overall interaction strength significantly. We used AlphaScreen as an orthogonal assay and obtained a 27-fold reduction in affinity upon ZnF6 deletion (ZnF456: IC_50_ = 79 ± 5 nM; ZnF45: IC_50_ = 2.15 ± 0.2 μM, see Fig. S6). Looking at the details of binding thermodynamics by ITC, with the assumption that ZnF4 and ZnF5 have equivalent thermodynamic contributions and that the contribution of ZnF6 is additive, we observe that ZnF4/5 provide a large enthalpic contribution (ΔH *=* -18.5 ± 0.5 kcal·M^-1^) whereas ZnF6 binding is not actually enthalpically favorable (ΔH *=* +0.9 ± 0.4 kcal·M^-1^). However, ZnF6 provides sizable entropic contribution to affinity (ΔS *=* -7.9 cal·M^-1^·°K^-1^) whereas ZnF4 and 5 pay a large entropic cost to bind DNA (ΔS *=* +46.9 cal·M^-1^·°K^-1^) (see Fig. S7). This observation is in accord with the more restricted conformation of ZnF6 in the solution ensemble, but higher level of dynamics retained by ZnF6 upon binding to DNA, translating to lower entropic cost.

**Figure 4.**
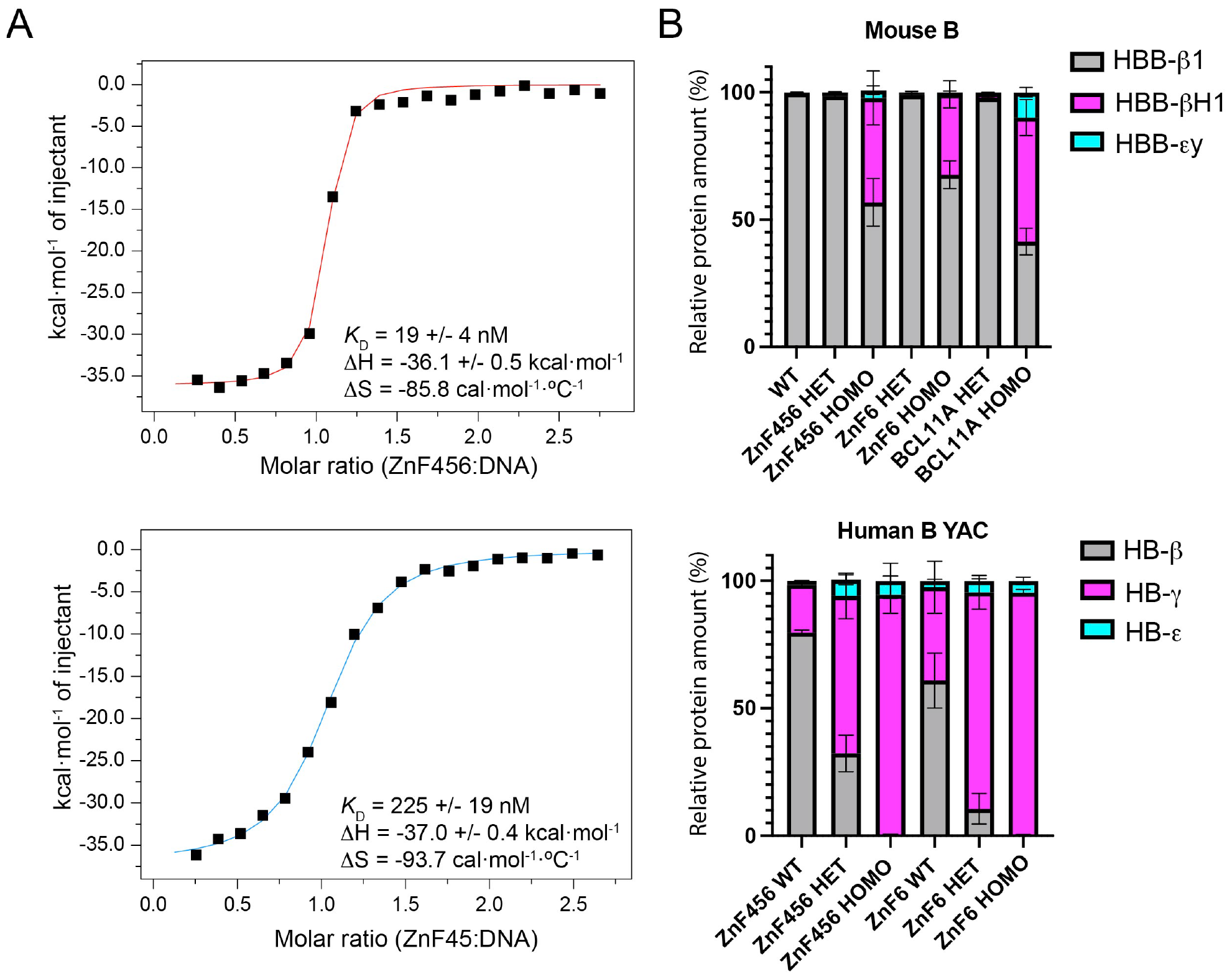
Identifying the special role of ZnF6 in DNA binding and fetal hemoglobin suppression. A) ITC measurement of ZnF456 (top) and ZnF45 (Δ6, bottom) binding to DNA. B) Quantification of β-like globin RNA transcripts in mice at FL stage. Genotypes include heterozygous and homozygous mice for ZnF456 deletion, ZnF6 deletion, and knockout of BCL11A gene. Small differences in maturation of embryos within a litter influence the precise level of γ-globin transcripts at E14.5d FL. Note, for example, the difference in γ-globin transcript levels in wild-type mice (ZnF456 WT and ZnF6 WT).

To delve deeper into the role of ZnF6, we excised it from the DNA-bound BCL11A complex and continued the MD simulation for an additional 1 microsecond. Beyond the anticipated loss of direct contacts due to its removal, we observed a consequential decrease in hydrogen bonds between ZnF4-5 and DNA. Specifically, the hydrogen bond occurrence between Asn753, Asn756, and Ser763 (part of ZnF5) and DNA diminished by 60%, 50%, and 59% respectively. These results suggest that ZnF6 enhances the overall hydrogen bond network, contributing to greater complex stability. This evidence underscores the distinct and critical function of ZnF6 in the DNA binding dynamics of BCL11A.

#### In vivo requirement for ZnF6 in globin repression

We next sought to establish an *in vivo* requirement for ZnF6 in the context of globin gene repression. Previously we have shown that the role of BCL11A in the control of globin switching can be revealed at the fetal liver (FL) stage of mouse development^24^. Through breeding of appropriate mouse strains, we can assess the fetal-to-adult transcriptional switch of both endogenous mouse globin genes and human globin genes contained in a yeast artificial chromosome (YAC) transgene. At the FL stage, mouse embryonic β-like globins, βH1 and εy, are silenced. The βH1 expression pattern serves as a surrogate for human γ-globin in the mouse. At the same time, the expression of transgenic human fetal (γ) and adult (β)-globin genes can be evaluated. Using CRISPR/Cas9 editing, we generated mice in which BCL11A, ZnF456 or ZnF6 alone were deleted from the endogenous mouse BCL11A locus (see Fig. S8). RNA transcripts of mouse and human globin genes in FL of mice embryos after 14 days of development (E14.5d) with various genotypes were quantitated. As shown in Fig. 4B, mice homozygous for loss of ZnF456 or ZnF6 exhibit markedly impaired silencing of βH1 and γ-globin expression. As a control, results for the endogenous mouse globins in the presence of knock-out (KO) of BCL11A are shown (Fig. 4B). The stronger effect of heterozygous loss of ZnF456 or ZnF6 on human γ-globin as compared with mouse βH1 highlights an intrinsic difference in the sensitivity of individual globin genes to loss of BCL11A function. However, the YAC transgene data is difficult to interpret due to apparent discrepancies in mice development stage between ZnF456 KO and ZnF6 KO groups.

The data for endogenous murine globins is clearer. Homozygous deletion of the entire BCL11A gene leads to a strong loss of fetal globin transcriptional silencing, with βH1 levels going back up to about 60% of the total globin mix (Fig. 4B top). Homozygous deletion of ZnF456 results in 45% βH1, and homozygous deletion of ZnF6 alone results in 35% βH1. The numbers can be seen as the transcriptional silencing power associated with each domain. In conclusion, the data provide *in vivo* evidence for the critical requirement for ZnF6 in transcriptional silencing, likely due to its distinct behavior in terms of conformation, dynamics, entropy, and ultimately DNA binding, and despite its lack of specific contacts with DNA.

## Discussion

In this study, we provide a detailed structural and dynamic analysis of the zinc finger-repeat region ZnF456 within BCL11A. We present a crystallographic structure of ZnF456 in complex with a 19-base pair DNA oligonucleotide that matches the distal promoter region of γ-globin. A longer DNA sequence was utilized to ensure an accurate conformation of ZnF6. Corroborating prior findings^17^ and supported by our NMR observations, ZnF6 interacts unspecifically with the minor groove of DNA, primarily contacting the DNA backbone.

The naturally occurring single-base substitutions that lead to increased HbF levels in adults in the HPFH syndrome can be rationalized from the DNA-bound structure. There are independent G-A mutations at position -117 that raise HbF levels from *ca*. 1% to 10-20%^15^. Notably, this is the only nucleotide that makes two specific contacts from the base to the protein. Asn756 and Ser783 have their OH groups 3.0 and 4.0 Å away from the oxygen atom of the guanine base, respectively. It is therefore understandable that G-A substitution would disrupt those contacts by replacing the hydrogen bond acceptor with a hydrogen bond donor at this position. At position -114, C-T and C-G mutations have been reported to elevate HbF levels to 5-15% and C-A seems to lead to lower HbF at *ca*. 3%^13,14^. Here, Arg787 makes two hydrogen bonds to the guanine in the complementary strand. Substitution for an adenosine or a cytosine would disrupt at least one of those hydrogen bonds, in line with a significantly impaired BCL11A binding affinity^13^. Substitution for a thymine could retain a hydrogen bond through the oxygen atom, although the resulting effect is more difficult to predict. The other nucleotides all contribute one specific contact with ZnF456, at the exception of -112 and -116 (although it could be involved in a contact with Gln781 as well), in line with their poor reported role in BCL11A binding.

Conversely, some aspects of the binding process of BCL11A to DNA are not recapitulated by the crystal structure alone. Meticulous examination of the apo solution structure and the binding thermodynamics uncovered a distinctive role for ZnF6, which significantly enhances the binding entropy and overall affinity of BCL11A for DNA. Repeats of three zinc finger domains are common in transcription factor and other nucleotide binding proteins. They provide an efficient way to create affinity and specificity for DNA with each ZnF typically recognizing three bases. This is true in the most general case when ZnF domains bind to the major groove of DNA^25^. However, binding to the minor groove has been reported as well like in GAGA^26^ and I-TevI^27^ and it is the case also for ZnF6 of BCL11A. This could be a feature enhancing affinity for DNA without extending the recognition site beyond the minimal promoter site.

Targeting transcription factors is often considered the holy grail in drug discovery. The targeting effort will benefit considerably from detailed structural information, including their DNA-free conformation and binding process. We determined the solution structure of apo ZnF456 using NMR and quantified the degree of conformational correlation between domains. Notably, zinc finger 6 is in a conformation that would clash with DNA and is correlated to ZnF5 to a higher degree than ZnF4. We show using NMR relaxation that all zinc fingers are well-folded and rigid in the absence of DNA. ZnF4 and 5 see their ps-ns dynamics lowered by DNA binding, such that they would bind tightly in a well-defined conformation at the cost of entropic loss. ZnF6, however, retains some of its flexibility both in the ps-ns and μs-ms time regimes even in the presence of DNA, explaining previous observations that it binds to the minor groove of the DNA helix and contributes less to the binding affinity^13,17^. However, the other side of the coin is that ZnF6 has positive entropic contribution to binding DNA and therefore participates in the affinity by a factor of 16. This specific role of ZnF6 is reflected in its *in vivo* requirement in globin gene switching (Fig. 4B).

In conclusion, we provide unprecedented detail into the structural analysis of BCL11A ZnF456 region and its mode of interaction with DNA. We suggest that these new findings may be leveraged for rational development of small molecules targeting binding of BCL11A to DNA that could be developed further in treatment of sickle cell disease and β-thalassemia. One of the findings here is that the apo-state of BCL11A is not compatible with DNA binding and must undergo a substantial structural change to engage DNA. The cost for this structural rearrangement is paid by favorable enthalpic contributions of ZnF4 and 5 and positive entropic contribution from ZnF6 to DNA binding. With such structural data, one might target transcriptional repression at different stages. Preventing BCL11A from binding to the DNA would be a preferred therapeutic route, however directly competing with DNA using a small molecule seems overly challenging. One plausible route is to find antibodies or nanobodies^28^ binding BCL11A and inducing protein degradation (PROTAC approach). Alternatively, small molecules could achieve the same results or stabilize the apo conformation of BCL11A preventing it from binding to DNA.

## Author contributions

Performed experiments: T.V., W.K., M.Y., A.J., Z.-Y.J.S., Y.F., K.Z., D.S., H.-S.S., S.D.P. Wrote initial draft: T.V. Commented and edited manuscript: all authors. Supervision: S.H.O., H.A.

## Acknowledgements

X-ray diffraction data were collected at the Northeastern Collaborative Access Team beamlines, which are funded by the National Institute of General Medical Sciences from the National Institutes of Health (P30 GM124165) and which used resources of the Advanced Photon Source, a U.S. Department of Energy (DOE) Office of Science User Facility operated for the DOE Office of Science by Argonne National Laboratory under Contract No. DE-AC02-06CH11357. Maintenance of the NMR instruments used for this research was supported by NIH grant No. EB002026. S.D.P. acknowledges funding from the Linde Family Foundation and the Doris Duke Charitable Foundation. H.A. acknowledges funding from the Claudia Adams Barr Program for Innovative Cancer Research and the R01 grant GM136859 from NIGMS. S.H.O. is an Investigator of the Howard Hughes Medical Institute and acknowledges partial support from NIH HL032259.

## Material and Methods

### Protein expression and purification

A construct of human BCL11A covering residues 730-835 in the pET28 was expressed in *E. coli* BL21 (DE3) in TB medium in the presence of 50 mg/ml of kanamycin. Cells were grown at 37°C to an OD of 0.6, induced overnight at 17°C with 500 μM isopropyl-1-thio-D-galactopyranoside, collected by centrifugation, and stored at -80°C. Cell pellets were microfluidized at 15,000 psi in buffer A (25 mM HEPES pH=7.5, 500 mM NaCl, 5% glycerol, 5 μM Zinc acetate, 20 mM imidazole, and 7 mM 2-mercapto-ethanol) and the resulting lysate was centrifuged at 13,000 rpm for 30 min. Ni-NTA beads (Qiagen) were mixed with lysate supernatant for 30 min and washed with buffer A and eluted with buffer C (25 mM HEPES pH=7.5, 500 mM NaCl, 5% glycerol, 5 μM Zinc acetate, 250 mM imidazole, and 7 mM 2-mercapto-ethanol). 3C protease was incubated with the sample overnight in the cold room. The sample was gel-filtered through an Superdex-75 16/600 column (GE healthcare) in buffer D (20 mM HEPES pH=7.5, 500 mM NaCl, 5% glycerol, 5 μM Zinc acetate, and 1 mM TCEP (Tris(2-carboxyethyl)phosphine)). Fractions were pooled, concentrated, and stored at -80°C. These fractions were used for crystallization.

For NMR and ITC, constructs encoding ZnF456 (residues 737-835) and ZnF45 (residues 737-795) were cloned into a pET28a vector containing a N-terminal His-SUMO tag for expression in *E. coli* in M9 minimal medium supplemented with ^15^NH4Cl and ^13^C-glucose. Proteins were purified as previously reported^28^.

### NMR spectroscopy

If not otherwise specified, NMR experiments were conducted on an Agilent DD2 spectrometer operating at 700 MHz, equipped with a triple-channel ^1^H, ^13^C, ^15^N cryogenically cooled probe. Samples of 200 μM ^15^N/^13^C-labeled BCL11A in PBS buffer pH=7.4 or pH=6.0, 1 mM dithiothreitol (DTT) and 10% v/v ^2^H_2_O were used for backbone assignments. Samples of approx. 100 μM ^15^N-labeled BCL11A (with or without 1.1:1 molar equivalent of 19-mer DNA) in PBS buffer pH=6.0, 1 mM dithiothreitol (DTT), 5 mM MgCl_2_ and 10% v/v ^2^H_2_O were used for relaxation studies. Experimental temperature was set to 25°C. All data were processed using NmrPipe^29^ and analyzed using CARA^30^ (for assignment) or CCPNmr Analysis^31^ (for relaxation).

#### Resonance assignment

This set of experiments is performed to obtain resonance assignments of the ZnF456 of BCL11A (correlating NMR signals to their respective amino acid nuclei). DNA-free BCL11A was assigned using a standard set of triple resonance backbone experiments: HNCA, HN(CO)CA, HNCO, HN(CA)CO, HN(CA)CB, CBCA(CO)NH for pH=7.4 sample, and HNCA, HN(CO)CA, HNCO, HN(CA)CB for pH=6.0 sample. Aliphatic side chains were assigned using (H)CC(CO)NH, H(CCCO)NH TOCSY experiments employing FLOPSY-16 mixing^32^. Aromatic side chains were assigned using (HB)CB(CGCD)HD and (HB)CB(CGCDCE)HE COSY experiments^33^. 3D experiments were recorded using non-uniform sampling(NUS), where Poisson-Gap sampling was used to select 10% of the Nyquist grid^34^ and the NUS data was reconstructed using the hmsIST protocol^35^. All residues could be assigned except for two N-terminal residues and the two proline residues at pH=6.0. However, ^15^N-^1^H HSQC peaks from residues K752, C754, S755, A780, S782, S783, S810, and Y812 were missing at pH=7.4. 82% of side chain proton resonances were assigned. Chemical shift assignments have been deposited in the BMRB under accession number 52029. Assignments were transferred to the DNA-bound BCL11A by visual inspection of the HSQC spectra and connectivities checked in an HNCA spectrum, 20 assignments could not be unambiguously transferred and hence were removed from further analysis.

Combined chemical shift perturbations were calculated as [(Δδ^1^H)^2^+(0.102·Δδ^15^N)^2^]^1/2^. Standard deviation to the mean was calculated excluding outliers with values higher than 3xSDM according to previous procedure^36^.

#### NMR restraints and structure calculation

This describes how distance, angular and orientational restraints are obtained and used for determination of the solution structure of DNA-free ZnF456. H_N_, N, CO, Cα, Cβ and Hα chemical shifts were used to derive dihedral angle values using the software TALOS-N^37. 15^N-, ^13^C- and aromatic-TROSY-edited NOESY spectra were collected using linear sampling, manually assigned and upper distance restraints were generated by CCPNmr Analysis^31^. H_N_-N RDCs were measured using C8E5:octanol liquid crystalline alignment medium^38^ (5 wt % C8E5:water and 0.87 molar ratio C8E5:octanol). The achieved D_2_O splitting was 12 Hz. Alignment tensors were calculated for each ZnF using the software REDCAT^39^ (Table S3). Upper distance hydrogen bond restraints (for residues forming α-helices) and Zn^2+^ restraints were set to standard values. All restraints were used in the software CYANA^40^, starting from 200 random structures with 50,000 simulated annealing steps. The 20 conformers with the lowest final CYANA target function values were further subjected to restrained energy minimization in explicit solvent using AMBER12 and the all atom force field ff99SB^41^. The structures were immersed in an octahedric box using the TIP3P water model, with a thickness of 10 Å. A total of 5 chlorine atoms were also included to neutralize charge. The simulation was performed under periodic boundary conditions and the particle-mesh Ewald approach was used to account for the electrostatic interactions. After energy minimization (2000 cycles) to regularize the CYANA structures, the temperature of the system was rapidly increased to 1000 K over 20 ps and then slowly cooled to 0 K over 250 ps. The distance and torsion angle constraints were applied with force constants of 32 kcal mol^−1^ Å^−2^ and 50 kcal mol^−1^ rad^−2^, respectively. These structures were inspected for any local regions of high restraint violation. The quality of the final ensemble was assessed using the PDB validation tool (Table S2) and submitted to the PDB under accession code 8THO.

#### Dynamics measurements by NMR

We used NMR based relaxation experiments to measure dynamics in the BCL11A protein in the free form and when bound to DNA. Nuclear spin relaxation (R_1_ and R_2_) and were measured with standard pulse sequences. R_2_ relaxation employed Carr-Purcell-Meiboom-Gill pulse trains with varied delays and corresponding temperature compensation blocks. Relaxation delays were set as follows (in ms):

- R_1_ DNA-free: 10, 100, 200, 400, 600, 900
- R_2_ DNA-free: 10, 30, 50, 110, 150
- R_1_ DNA-bound: 10, 200, 400, 800
- R_2_ DNA-bound: 10, 30, 50, 70, 110

Steady-state heteronuclear nuclear Overhauser effect (hetNOE) were measured on a Bruker Avance III spectrometer operating at 750 MHz equipped with a triple-channel TCI ^1^H, ^13^C, ^15^N cryogenically cooled probe. ^1^H saturation was achieved with a train of 180° pulses at 11.25 kHz power for a delay of 6 s (> 4·T1^max^), recycling delays of 10 s were employed.

Carr-Purcell-Meiboom-Gill (CPMG) relaxation dispersion experiments were conducted on an Agilent DD2 spectrometer operating at 600 MHz, equipped with a triple-channel ^1^H, ^13^C, ^15^N cryogenically cooled probe. T_2_ relaxation delay was set to 40 ms CPMG frequencies were modulated in the following order (in Hz): 0, 1000, 50, 850, 100, 650, 500, 250, 400, 100, 50, 850, 400, 1000. Dispersion curves were analyzed with the program Relax^42^.

### X-ray crystallography

#### Crystallization

Samples of 400 μM ZnF456 and 600 μM 12-bp (GCTTGACCAATGC with 1-bp overhang) or 19-bp (GCTTGACCAATGCGGTCGC with 1-bp overhang) duplex DNA were co-crystallized in 30% PEG6K and 0.1 M Na Citrate pH=5.0 by sitting-drop vapor diffusion at 4°C using a combination of Formulatrix NT8 and ArtRobbins Phoenix liquid handlers and visualized using a Formulatrix RockImager. Large, single crystals were transferred briefly into crystallization buffer containing 25% glycerol prior to flash-freezing in liquid nitrogen and shipped to the synchrotron for data collection.

#### Data collection and structure determination

Diffraction data were collected at beamline 24ID-C of the NE-CAT at the Advanced Photon Source (Argonne National Laboratory). Data sets were processed using XDS^43^. Structures were solved by molecular replacement using the program Phaser^44^ and the search model (PDB entry 1MEY). Iterative manual model building and refinement using Phenix^45^ and Coot^46^ led to a model with excellent statistics, shown in Table S1. This work used SBGrid compiled software^47^. Final coordinates were deposited in the PDB under accession number 6U9Q for the 12-bp DNA complex and 8TLO for the 19-bp DNA complex.

### Molecular dynamics simulations

Structures of BCL11A ZnF456 and DNA (5’-ATATTGGTCAAGG-3’) were retrieved from the PDB (6KI6)^17^. Crystallographic water molecules and other non-relevant biomolecular moieties were removed. The missing loop residues 792-797 were modelled using Modeller4^48^. Additional DNA sequence was appended to increase the length from 13 to 21 nucleotides and achieve two complete turns of helical B-DNA (5’-AAGGCTATATTGGTCAAGGCAAG-3’). All Zn ions were modelled using ZAFF methodology^49^ implemented in AMBER v216. BCL11A protein was parameterized using ff14SB forcefield^41^ and DNA was parametrized using Bsc0 forcefield^50^. CMAP modifications with environmental specific precise force field (ESFF1)^50^ were added over ff14SB forcefield to account for its high flexibility.

For binding simulations, BCL11A was placed 25 Å away parallel to the DNA strands. Restraints were placed on Arg787 to prevent BCL11A diffusing more than 30 Å away from G114 of DNA. No restraints were applied to attract BCL11A to DNA. The model was solvated in TIP4P-B water model^51^ with a clearance of 10 Å from the edges of the solvated cubic box. The complex was electro neutralized and brought to 0.15 mM ionic strength using Na^+^ and Cl^-^ ions. Periodic boundary conditions along with particle mesh Ewald utilizing 10 Å cutoff for long-range electrostatic interactions was deployed. Movement of bonded hydrogen atoms was restricted using the SHAKE algorithm. This enabled use of 2 fs time steps during simulation. Temperature and isotropic pressure regulation used Langevin dynamics thermostat and Berendsen barostat, respectively. The construct was energy minimized using a 5-step process (each step contained 3000 steps of steepest decent and 2000 steps of conjugate gradient), successively decreasing the restraints on protein and DNA from 100 kcal/mol to 0 kcal/mol. The construct was then heated to 300 K over 20 ps. Equilibration employed a 5-step NVT ensemble simulation process with successively decreasing restraints on protein and DNA from 100 kcal/mol to 0 kcal/mol. Each equilibration step was run for 100 ps. This was followed with 20 ns of unrestricted equilibration simulation in NPT conditions. Total, potential, and kinetic energy along with RMSD of individual biomolecular entities were monitored for stability. The production run of MD used NPT ensemble with snapshots saved every 20 ps. All MD was performed using pmemd^52,53^ in AMBER v21 on nVIDIA RTX A5000 GPUs.

### Isothermal titration calorimetry

ITC experiments were carried out on a Malvern Microcal ITC200. All experiments were carried out in a buffer that contained 20 mM HEPES pH=7.5, 150 mM NaCl, and 1 mM DTT, with 2% DMSO at 25 °C. The protein BCL11A was held at a concentration of 20 μM in the calorimetric cell. Double stranded DNA (5’-CTTGACCAAT-3’) at a stock concentration of 200 μM was titrated in 2.5 μL injections at 200 s intervals using stirring speed at 125 rpm. Resulting isotherm was fitted with a single site model to derive thermodynamic parameters of ΔH, ΔS, stoichiometry and *K*_d_.

### Alpha Screen

The assay was performed in 384-well plates where a constant concentration of biotinylated BCL11A ZnF456 or ZnF45 was incubated with varying concentrations of 14-bp double stranded DNA (5’-GCTTGACCAATGC-3’) for 30 min at room temperature. 10 µl of streptavidin donor beads and 10 µl of nickel chelate (Ni-NTA) acceptor beads (PerkinElmer) in assay buffer were added and incubated in dark for 1 hr. After centrifuge for 15 s at 161×g, the fluorescent signal was measured by a plate reader at 580 nm after excitation at 680 nM.

### Generation and assessment of globin transcripts in fetal mice liver

ZnF456 and ZnF6 deletion mutant alleles were generated as by-products of aberrant targeting in the process of knocking-in variant FKBP^F36V^ sequences into the BCL11A locus of C57/Bl mice^54^. The detailed strategy for generation of the knock-in allele, including oligonucleotide primers and sgRNA sequences, were previously reported^54^. A schematic representation of the BCL11A locus and the resultant ZnF456 and ZnF6 deletion alleles is summarized in Supplementary Fig. 8. In the ZnF456 deletion, 298 bp were removed, which encode residues 742-835 of BCL11A-XL. The deletion generates a frameshift, such that 7 residues (RPSLTPT) are added at the C-terminus of the truncated protein. The ZnF6 deletion allele lacks 110 bp, encompassing residues 803-835. The same genotyping primers were used for both alleles:

- BCL11A_Cterm_F1, 5’-AATCGCCTTTTGCCTCCTCA-3’
- BCL11A_Cterm_R1, 5’-GTTAGTCAAACCCAGACACCGT-3’

PCR products were: wild-type: 1042 bp; ZnF456 deletion: 744 bp; and ZnF6 deletion: 932 bp. Mice harboring a wild-type human β-globin gene containing YAC transgene (β-YAC)^55^ were bred with ZnF456 and ZnF6 deletion mice. Litters were sacrificed at E14.5 FL to analyze globin gene transcripts, as previously described^24^.

## References

1. Weatherall, D.J. (2010). The inherited diseases of hemoglobin are an emerging global health burden. Blood 115, 4331–4336. 10.1182/blood-2010-01-251348.

2. Sankaran, V.G., and Weiss, M.J. (2015). Anemia: progress in molecular mechanisms and therapies. Nat Med 21, 221–230. 10.1038/nm.3814.

3. Platt, O.S., Brambilla, D.J., Rosse, W.F., Milner, P.F., Castro, O., Steinberg, M.H., and Klug, P.P. (1994). Mortality in sickle cell disease. Life expectancy and risk factors for early death. N Engl J Med 330, 1639–1644. 10.1056/NEJM199406093302303.

4. Galanello, R., Sanna, S., Perseu, L., Sollaino, M.C., Satta, S., Lai, M.E., Barella, S., Uda, M., Usala, G., Abecasis, G.R., and Cao, A. (2009). Amelioration of Sardinian beta0 thalassemia by genetic modifiers. Blood 114, 3935–3937. 10.1182/blood-2009-04-217901.

5. Andrieu-Soler, C., and Soler, E. (2020). When basic science reaches into rational therapeutic design: from historical to novel leads for the treatment of beta-globinopathies. Curr Opin Hematol 27, 141–148. 10.1097/MOH.0000000000000577.

6. Uda, M., Galanello, R., Sanna, S., Lettre, G., Sankaran, V.G., Chen, W., Usala, G., Busonero, F., Maschio, A., Albai, G., et al. (2008). Genome-wide association study shows BCL11A associated with persistent fetal hemoglobin and amelioration of the phenotype of beta-thalassemia. Proc Natl Acad Sci U S A 105, 1620–1625. 10.1073/pnas.0711566105.

7. Menzel, S., Garner, C., Gut, I., Matsuda, F., Yamaguchi, M., Heath, S., Foglio, M., Zelenika, D., Boland, A., Rooks, H., et al. (2007). A QTL influencing F cell production maps to a gene encoding a zinc-finger protein on chromosome 2p15. Nat Genet 39, 1197–1199. 10.1038/ng2108.

8. Canver, M.C., Smith, E.C., Sher, F., Pinello, L., Sanjana, N.E., Shalem, O., Chen, D.D., Schupp, P.G., Vinjamur, D.S., Garcia, S.P., et al. (2015). BCL11A enhancer dissection by Cas9-mediated in situ saturating mutagenesis. Nature 527, 192–197. 10.1038/nature15521.

9. Traxler, E.A., Yao, Y., Wang, Y.D., Woodard, K.J., Kurita, R., Nakamura, Y., Hughes, J.R., Hardison, R.C., Blobel, G.A., Li, C., and Weiss, M.J. (2016). A genome-editing strategy to treat beta-hemoglobinopathies that recapitulates a mutation associated with a benign genetic condition. Nat Med 22, 987–990. 10.1038/nm.4170.

10. Sankaran, V.G., Menne, T.F., Xu, J., Akie, T.E., Lettre, G., Van Handel, B., Mikkola, H.K., Hirschhorn, J.N., Cantor, A.B., and Orkin, S.H. (2008). Human fetal hemoglobin expression is regulated by the developmental stage-specific repressor BCL11A. Science 322, 1839–1842. 10.1126/science.1165409.

11. Xu, J., Peng, C., Sankaran, V.G., Shao, Z., Esrick, E.B., Chong, B.G., Ippolito, G.C., Fujiwara, Y., Ebert, B.L., Tucker, P.W., and Orkin, S.H. (2011). Correction of sickle cell disease in adult mice by interference with fetal hemoglobin silencing. Science 334, 993–996. 10.1126/science.1211053.

12. Xu, J., Sankaran, V.G., Ni, M., Menne, T.F., Puram, R.V., Kim, W., and Orkin, S.H. (2010). Transcriptional silencing of {gamma}-globin by BCL11A involves long-range interactions and cooperation with SOX6. Genes Dev 24, 783–798. 10.1101/gad.1897310.

13. Liu, N., Hargreaves, V.V., Zhu, Q., Kurland, J.V., Hong, J., Kim, W., Sher, F., Macias-Trevino, C., Rogers, J.M., Kurita, R., et al. (2018). Direct Promoter Repression by BCL11A Controls the Fetal to Adult Hemoglobin Switch. Cell 173, 430–442 e417. 10.1016/j.cell.2018.03.016.

14. Collins, F.S., Metherall, J.E., Yamakawa, M., Pan, J., Weissman, S.M., and Forget, B.G. (1985). A point mutation in the A gamma-globin gene promoter in Greek hereditary persistence of fetal haemoglobin. Nature 313, 325–326. 10.1038/313325a0.

15. Gelinas, R., Endlich, B., Pfeiffer, C., Yagi, M., and Stamatoyannopoulos, G. (1985). G to A substitution in the distal CCAAT box of the A gamma-globin gene in Greek hereditary persistence of fetal haemoglobin. Nature 313, 323–325. 10.1038/313323a0.

16. Martyn, G.E., Wienert, B., Yang, L., Shah, M., Norton, L.J., Burdach, J., Kurita, R., Nakamura, Y., Pearson, R.C.M., Funnell, A.P.W., et al. (2018). Natural regulatory mutations elevate the fetal globin gene via disruption of BCL11A or ZBTB7A binding. Nat Genet 50, 498–503. 10.1038/s41588-018-0085-0.

17. Yang, Y., Xu, Z., He, C., Zhang, B., Shi, Y., and Li, F. (2019). Structural insights into the recognition of gamma-globin gene promoter by BCL11A. Cell Res 29, 960–963. 10.1038/s41422-019-0221-0.

18. Patel, A., Yang, P., Tinkham, M., Pradhan, M., Sun, M.A., Wang, Y., Hoang, D., Wolf, G., Horton, J.R., Zhang, X., et al. (2018). DNA Conformation Induces Adaptable Binding by Tandem Zinc Finger Proteins. Cell 173, 221–233 e212. 10.1016/j.cell.2018.02.058.

19. Park, S., Phukan, P.D., Zeeb, M., Martinez-Yamout, M.A., Dyson, H.J., and Wright, P.E. (2017). Structural Basis for Interaction of the Tandem Zinc Finger Domains of Human Muscleblind with Cognate RNA from Human Cardiac Troponin T. Biochemistry 56, 4154–4168. 10.1021/acs.biochem.7b00484.

20. Payne, M.A. (2011). Zinc finger structure-function in Ikaros Marvin A Payne. World J Biol Chem 2, 161–166. 10.4331/wjbc.v2.i6.161.

21. Henley, M.J., and Koehler, A.N. (2021). Advances in targeting ‘undruggable’ transcription factors with small molecules. Nat Rev Drug Discov 20, 669–688. 10.1038/s41573-021-00199-0.

22. Tochio, N., Umehara, T., Nakabayashi, K., Yoneyama, M., Tsuda, K., Shirouzu, M., Koshiba, S., Watanabe, S., Kigawa, T., Sasazuki, T., et al. (2015). Solution structures of the DNA-binding domains of immune-related zinc-finger protein ZFAT. J Struct Funct Genomics 16, 55–65. 10.1007/s10969-015-9196-3.

23. Jaremko, L., Jaremko, M., Ejchart, A., and Nowakowski, M. (2018). Fast evaluation of protein dynamics from deficient (15)N relaxation data. J Biomol NMR 70, 219–228. 10.1007/s10858-018-0176-3.

24. Sankaran, V.G., Xu, J., Ragoczy, T., Ippolito, G.C., Walkley, C.R., Maika, S.D., Fujiwara, Y., Ito, M., Groudine, M., Bender, M.A., et al. (2009). Developmental and species-divergent globin switching are driven by BCL11A. Nature 460, 1093–1097. 10.1038/nature08243.

25. Pavletich, N.P., and Pabo, C.O. (1991). Zinc finger-DNA recognition: crystal structure of a Zif268-DNA complex at 2.1 A. Science 252, 809–817. 10.1126/science.2028256.

26. Eom, K.S., Cheong, J.S., and Lee, S.J. (2016). Structural Analyses of Zinc Finger Domains for Specific Interactions with DNA. J Microbiol Biotechnol 26, 2019–2029. 10.4014/jmb.1609.09021.

27. Krishna, S.S., Majumdar, I., and Grishin, N.V. (2003). Structural classification of zinc fingers: survey and summary. Nucleic Acids Res 31, 532–550. 10.1093/nar/gkg161.

28. Yin, M., Izadi, M., Tenglin, K., Viennet, T., Zhai, L., Zheng, G., Arthanari, H., Dassama, L.M.K., and Orkin, S.H. (2023). Evolution of nanobodies specific for BCL11A. Proceedings of the National Academy of Sciences of the United States of America 120, e2218959120. 10.1073/pnas.2218959120.

29. Delaglio, F., Grzesiek, S., Vuister, G.W., Zhu, G., Pfeifer, J., and Bax, A. (1995). NMRPipe: a multidimensional spectral processing system based on UNIX pipes. J Biomol NMR 6, 277–293. 10.1007/bf00197809.

30. Keller, R.L.J. (2004). The computer aided resonance assignment tutorial. In (CANTINA Verlag).

31. Vranken, W.F., Boucher, W., Stevens, T.J., Fogh, R.H., Pajon, A., Llinas, M., Ulrich, E.L., Markley, J.L., Ionides, J., and Laue, E.D. (2005). The CCPN data model for NMR spectroscopy: development of a software pipeline. Proteins 59, 687–696. 10.1002/prot.20449.

32. Kovacs, H., and Gossert, A. (2014). Improved NMR experiments with (1)(3)C-isotropic mixing for assignment of aromatic and aliphatic side chains in labeled proteins. J Biomol NMR 58, 101–112. 10.1007/s10858-013-9808-9.

33. Yamazaki, T., Forman-Kay, J.D., and Kay, L.E. (2002). Two-dimensional NMR experiments for correlating carbon-13.beta. and proton.delta./.epsilon. chemical shifts of aromatic residues in 13C-labeled proteins via scalar couplings. Journal of the American Chemical Society 115, 11054–11055. 10.1021/ja00076a099.

34. Hyberts, S.G., Takeuchi, K., and Wagner, G. (2010). Poisson-gap sampling and forward maximum entropy reconstruction for enhancing the resolution and sensitivity of protein NMR data. J Am Chem Soc 132, 2145–2147. 10.1021/ja908004w.

35. Hyberts, S.G., Milbradt, A.G., Wagner, A.B., Arthanari, H., and Wagner, G. (2012). Application of iterative soft thresholding for fast reconstruction of NMR data non-uniformly sampled with multidimensional Poisson Gap scheduling. J Biomol NMR 52, 315–327. 10.1007/s10858-012-9611-z.

36. Schumann, F.H., Riepl, H., Maurer, T., Gronwald, W., Neidig, K.P., and Kalbitzer, H.R. (2007). Combined chemical shift changes and amino acid specific chemical shift mapping of protein-protein interactions. J Biomol NMR 39, 275–289. 10.1007/s10858-007-9197-z.

37. Shen, Y., and Bax, A. (2013). Protein backbone and sidechain torsion angles predicted from NMR chemical shifts using artificial neural networks. J Biomol NMR 56, 227–241. 10.1007/s10858-013-9741-y.

38. Rückert, M., and Otting, G. (2000). Alignment of Biological Macromolecules in Novel Nonionic Liquid Crystalline Media for NMR Experiments. Journal of the American Chemical Society 122, 7793–7797. 10.1021/ja001068h.

39. Valafar, H., and Prestegard, J.H. (2004). REDCAT: a residual dipolar coupling analysis tool. J Magn Reson 167, 228–241. 10.1016/j.jmr.2003.12.012.

40. Guntert, P., and Buchner, L. (2015). Combined automated NOE assignment and structure calculation with CYANA. J Biomol NMR 62, 453–471. 10.1007/s10858-015-9924-9.

41. Maier, J.A., Martinez, C., Kasavajhala, K., Wickstrom, L., Hauser, K.E., and Simmerling, C. (2015). ff14SB: Improving the Accuracy of Protein Side Chain and Backbone Parameters from ff99SB. J Chem Theory Comput 11, 3696–3713. 10.1021/acs.jctc.5b00255.

42. Morin, S., Linnet, T.E., Lescanne, M., Schanda, P., Thompson, G.S., Tollinger, M., Teilum, K., Gagne, S., Marion, D., Griesinger, C., et al. (2014). Relax: the analysis of biomolecular kinetics and thermodynamics using NMR relaxation dispersion data. Bioinformatics 30, 2219–2220. 10.1093/bioinformatics/btu166.

43. Kabsch, W. (2010). Integration, scaling, space-group assignment and post-refinement. Acta Crystallogr D Biol Crystallogr 66, 133–144. 10.1107/S0907444909047374.

44. McCoy, A.J., Grosse-Kunstleve, R.W., Adams, P.D., Winn, M.D., Storoni, L.C., and Read, R.J. (2007). Phaser crystallographic software. J Appl Crystallogr 40, 658–674. 10.1107/S0021889807021206.

45. Adams, P.D., Afonine, P.V., Bunkoczi, G., Chen, V.B., Davis, I.W., Echols, N., Headd, J.J., Hung, L.W., Kapral, G.J., Grosse-Kunstleve, R.W., et al. (2010). PHENIX: a comprehensive Python-based system for macromolecular structure solution. Acta Crystallogr D Biol Crystallogr 66, 213–221. 10.1107/S0907444909052925.

46. Emsley, P., and Cowtan, K. (2004). Coot: model-building tools for molecular graphics. Acta Crystallogr D Biol Crystallogr 60, 2126–2132. 10.1107/S0907444904019158.

47. Morin, A., Eisenbraun, B., Key, J., Sanschagrin, P.C., Timony, M.A., Ottaviano, M., and Sliz, P. (2013). Collaboration gets the most out of software. Elife 2, e01456. 10.7554/eLife.01456.

48. Webb, B., and Sali, A. (2016). Comparative Protein Structure Modeling Using MODELLER. Curr Protoc Bioinformatics 54, 5 6 1-5 6 37. 10.1002/cpbi.3.

49. Peters, M.B., Yang, Y., Wang, B., Fusti-Molnar, L., Weaver, M.N., and Merz, K.M., Jr. (2010). Structural Survey of Zinc Containing Proteins and the Development of the Zinc AMBER Force Field (ZAFF). J Chem Theory Comput 6, 2935–2947. 10.1021/ct1002626.

50. Love, O., Galindo-Murillo, R., Zgarbova, M., Sponer, J., Jurecka, P., and Cheatham, T.E., 3rd (2023). Assessing the Current State of Amber Force Field Modifications for DNA horizontal line 2023 Edition. J Chem Theory Comput 19, 4299–4307. 10.1021/acs.jctc.3c00233.

51. Mu, J., Pan, Z., and Chen, H.F. (2021). Balanced Solvent Model for Intrinsically Disordered and Ordered Proteins. J Chem Inf Model 61, 5141–5151. 10.1021/acs.jcim.1c00407.

52. Gotz, A.W., Williamson, M.J., Xu, D., Poole, D., Le Grand, S., and Walker, R.C. (2012). Routine Microsecond Molecular Dynamics Simulations with AMBER on GPUs. 1. Generalized Born. J Chem Theory Comput 8, 1542–1555. 10.1021/ct200909j.

53. Salomon-Ferrer, R., Gotz, A.W., Poole, D., Le Grand, S., and Walker, R.C. (2013). Routine Microsecond Molecular Dynamics Simulations with AMBER on GPUs. 2. Explicit Solvent Particle Mesh Ewald. J Chem Theory Comput 9, 3878–3888. 10.1021/ct400314y.

54. Mehta, S., Buyanbat, A., Kai, Y., Karayel, O., Goldman, S.R., Seruggia, D., Zhang, K., Fujiwara, Y., Donovan, K.A., Zhu, Q., et al. (2022). Temporal resolution of gene derepression and proteome changes upon PROTAC-mediated degradation of BCL11A protein in erythroid cells. Cell Chem Biol 29, 1273–1287 e1278. 10.1016/j.chembiol.2022.06.007.

55. Porcu, S., Kitamura, M., Witkowska, E., Zhang, Z., Mutero, A., Lin, C., Chang, J., and Gaensler, K.M.L. (1997). The Human β Globin Locus Introduced by YAC Transfer Exhibits a Specific and Reproducible Pattern of Developmental Regulation in Transgenic Mice. Blood 90, 4602–4609. 10.1182/blood.V90.11.4602.

