## Supplemental Information for "Structural Insights into the DNA-Binding Mechanism of BCL11A: The Integral Role of ZnF6"

### Supplementary Tables

**Table S1. Crystallization conditions and data collection and refinement statistics for complex crystal structure.** Values in parenthesis are for highest resolution shell.

| **Protein** | BCL11A ZnF456 |
|---|---|
| **Ligand** | GCTTGACCAATGC |
| **RSCB accession code** | 6U9Q |
| **Data collection** | |
| Space group | C2 |
| Cell dimensions a, b, c (Å) | 101.00, 60.16, 75.79 |
| Cell dimensions a, b, g (°) | 90.00, 129.73, 90.00 |
| Resolution (Å) | 47.46 – 1.83 (1.89 – 1.83) |
| $R_{pim}$ | 0.026 (0.579) |
| Redundancy | 3.3 (3.4) |
| Completeness (%) | 97.28 (97.99) |
| σ | 15.79 (1.21) |
| **Structure solution** | |
| PDB entry for molecular replacement | 1MEY |
| **Refinement** | |
| No. reflections | 30108 (3022) |
| $R_{work}$ | 0.1758 (0.3418) |
| $R_{free}$ | 0.1953 (0.3570) |
| No. atoms | 1329 |
| Macromolecules | 1206 |
| Ligand/ions | 14 |
| Water | 109 |
| B-factors | 52.4 |
| Macromolecules | 52.3 |
| Ligand/ions | 72.2 |
| Water | 51.4 |
| Bond length r.m.s.d (Å) | 0.008 |
| Bond angle r.m.s.d. (°) | 1.13 |
| **Ramachandran** | |
| Preferred (%) | 97.70 |
| Allowed (%) | 2.30 |
| Not allowed (%) | 0.00 |

**Table S2. Solution structure restraints and statistics for DNA-free ZnF456.**

| | |
|---|---|
| ***NMR distance constraints*** | |
| Total NOEs | 1352 |
| Intra-residue | 741 |
| Inter-residue | 611 |
| Sequential (\|i-j\|=1) | 333 |
| Medium-range (\|i-j\|=1) | 143 |
| Long-range (\|i-j\|=1) | 135 |
| ***Dihedral angle constraints*** | 126 |
| ***Hydrogen bond constraints*** | 32 |
| ***Zn ion distance constraints*** | 24 |
| ***RDC constraints*** | 50 |
| ***Violations (mean ± s.d.)*** | |
| Distance constraints (Å) | 0.095 ± 0.004 |
| Dihedral angles constraints (°) | 4.80 ± 0.54 |
| RDC constraints (Hz) | 0.91 ± 0.24 |
| ***Deviations from idealized geometry*** | |
| Bond length (Å) | 0.0 |
| Bond angles (°) | 0.0 |
| ***Ramachandran plot analysis*** | |
| Favored (%) | 80 ± 3 |
| Allowed (%) | 15 ± 3 |
| Outliers (%) | 5 ± 2 |
| ***RDC tensor Q-factor (%)*** | |
| ZnF4 (aa 5-30) | 19.5 ± 3.3 |
| ZnF5 (aa 35-57) | 12.4 ± 1.3 |
| ZnF6 (aa 65-88) | 15.7 ± 0.8 |
| ***Average pairwise rmsd (Å, backbone / heavy atoms)*** | |
| ZnF4 (aa 5-30) | 0.81 ± 0.28 / 1.49 ± 0.42 |
| ZnF5 (aa 35-57) | 0.45 ± 0.14 / 1.12 ± 0.24 |
| ZnF6 (aa 65-88) | 0.39 ± 0.12 / 1.00 ± 0.22 |
| ZnF456 (aa 1-100) | 4.06 ± 0.86 / 4.46 ± 0.86 |

**Table S3. RDC tensor fitting results and statistics.**

| | Magnitude, Da | Rhombicity, R | Q-factor (%) |
|---|---|---|---|
| ZnF4 (aa 5-30) | 13.5 | 0.32 | 12 |
| ZnF5 (aa 35-57) | 20.8 | 0.29 | 8 |
| ZnF6 (aa 65-88) | 16.0 | 0.42 | 13 |
| ZnF4-ZnF5 | 16.2 | 0.47 | 65 |
| ZnF5-ZnF6 | 16.9 | 0.39 | 62 |
| ZnF4-ZnF6 | 9.5 | 0.08 | 62 |
| ZnF456 | 10.8 | 0.49 | 81 |

### Supplementary Figures

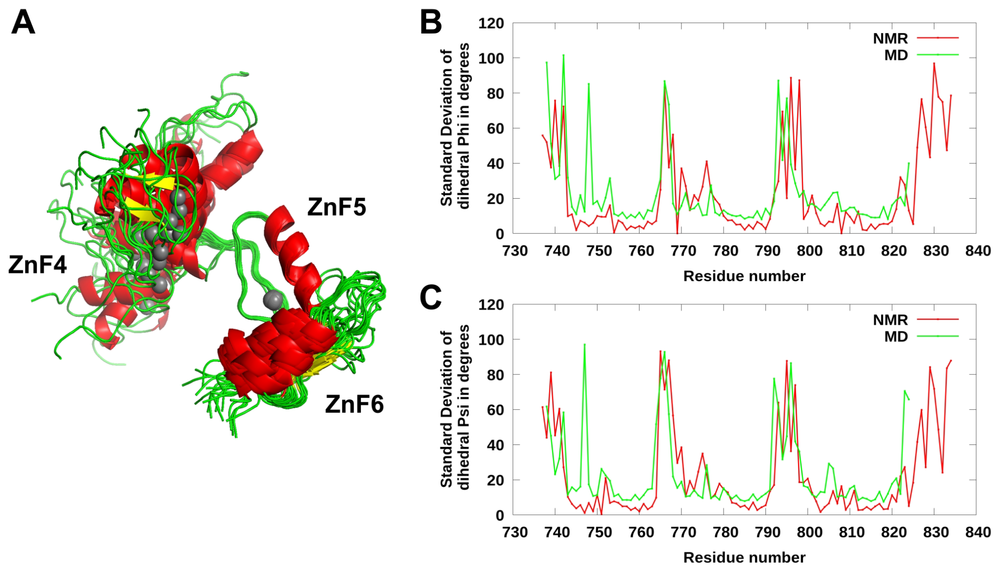

**Supplementary Figure S1. Molecular dynamics simulation of apo BCL11A.** A) MD trajectory ensemble aligned on ZnF5. B-C) Comparison of residue-specific φ (B) and ψ (C) dihedral angles in the NMR (red) and MD (green) ensembles.

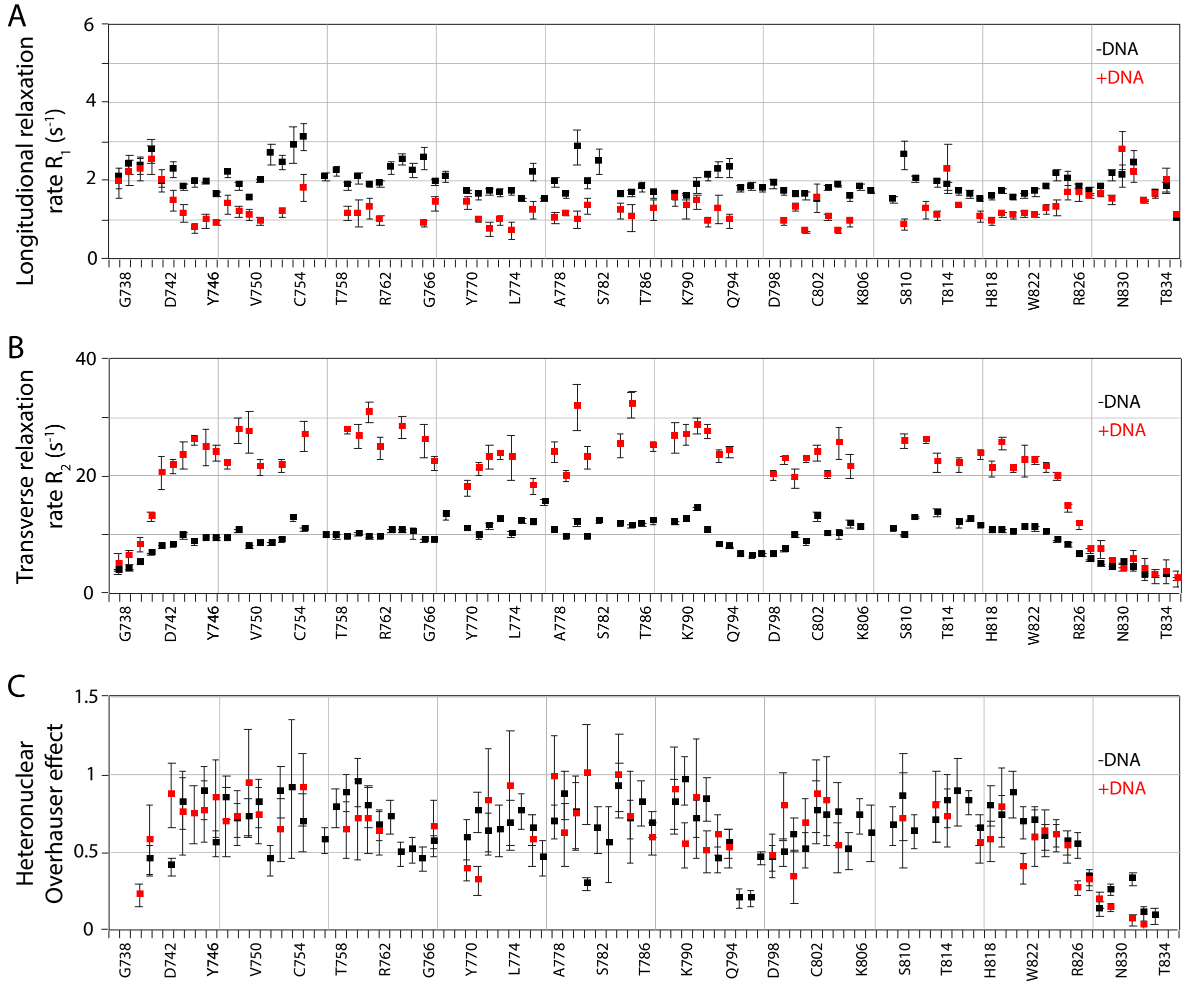

**Supplementary Figure S2. Picosecond-nanosecond motions in BCL11A free and bound to DNA.** A) $^{15}$N longitudinal ($R_1$) relaxation rates for each residue of ZnF456 in the absence (black) and presence (red) of DNA. B) $^{15}$N transverse ($R_2$) relaxation rates for each residue of ZnF456 in the absence (black) and presence (red) of DNA. C) {$^{1}$H}$^{15}$N heteronuclear Overhauser effect for each residue of ZnF456 in the absence (black) and presence (red) of DNA.

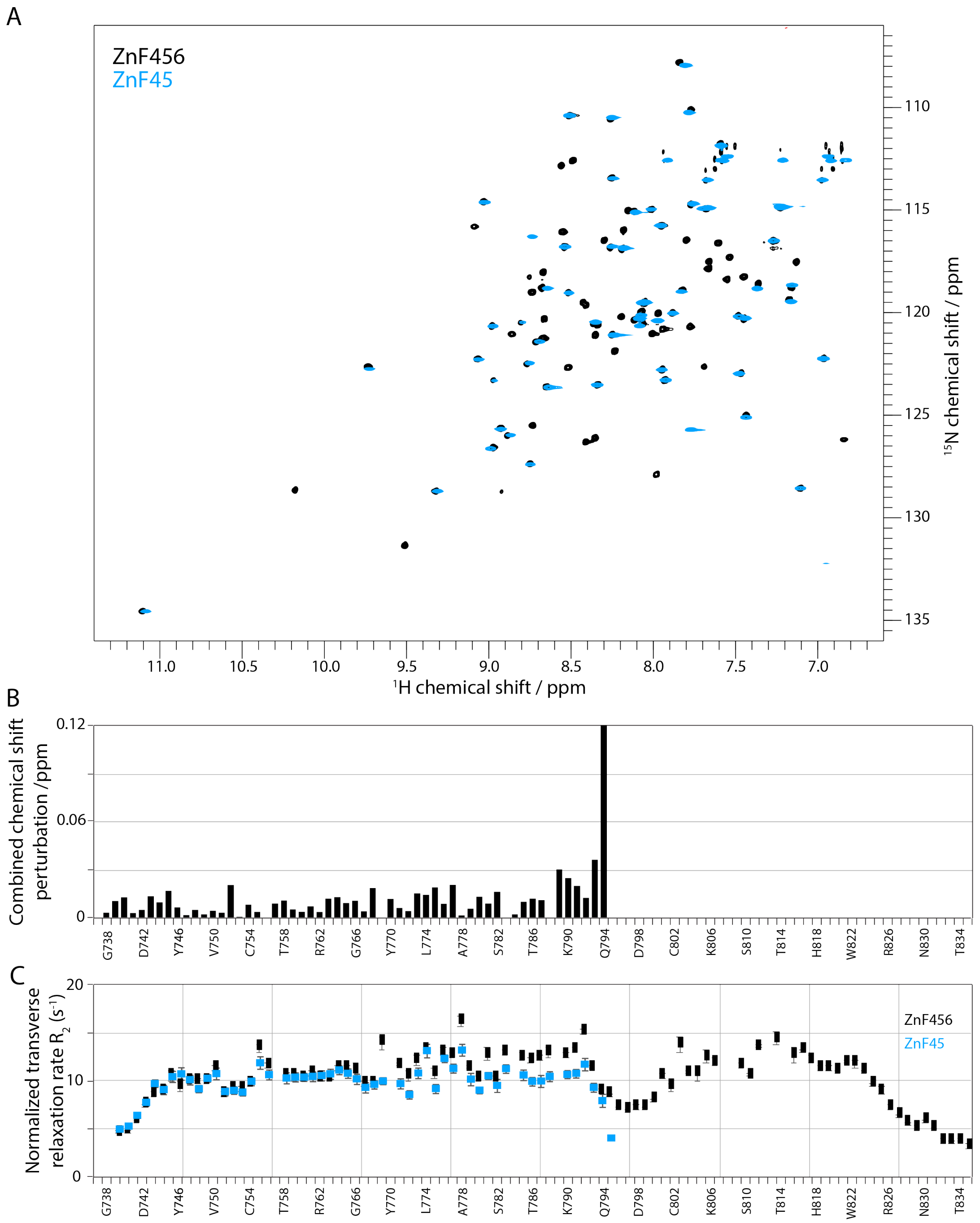

**Supplementary Figure S3. Truncation of ZnF6 dynamically affects ZnF5 but not ZnF4.** A) $^{15}$N-$^{1}$H HSQC spectrum of ZnF456 (black) and ZnF45 (cyan). B) Corresponding combined chemical shift perturbation plotted against the amino acid sequence of ZnF456. C) $^{15}$N transverse ($R_2$) relaxation rates for each residue of ZnF456 (black) or ZnF45 (cyan). Note that R2 rates have been normalized to the first residue to account for the different correlation times of the different molecular weight constructs.

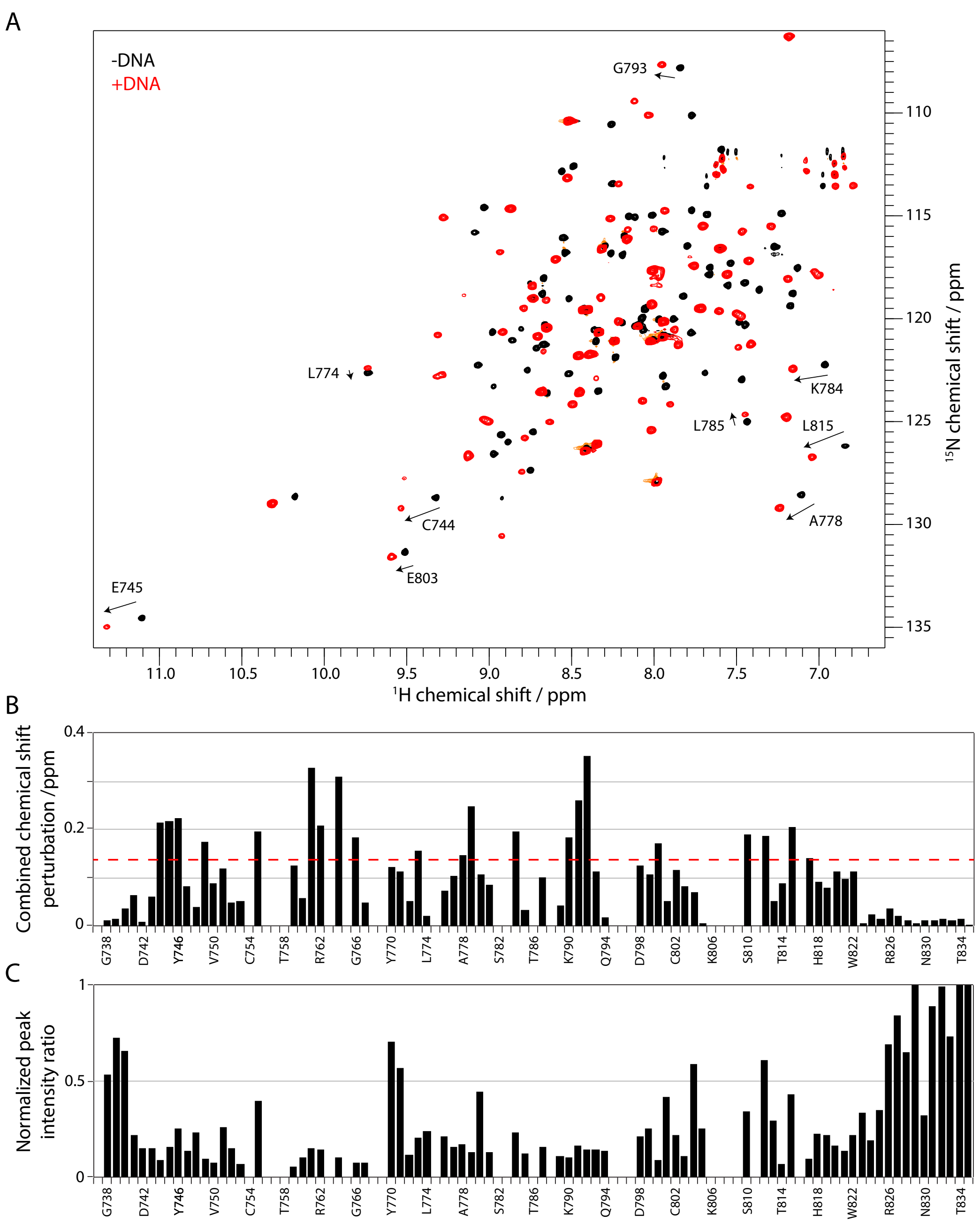

**Supplementary Figure S4. NMR identifies the region of BCL11A that bind DNA.** A) $^{15}N$-$^{1}H$ HSQC spectrum of ZnF456 in the absence (black) and presence (red) of a DNA. Select residue assignment are shown and an arrow indicates how they shift upon addition of DNA. B) Corresponding combined chemical shift perturbation plotted against the amino acid sequence of ZnF456. C) Corresponding normalized peak intensity ratio ($I_{+DNA}$ / $I_{-DNA}$) plotted against the amino acid sequence of ZnF456. Intensity of the C-terminal peak was normalized to 1.

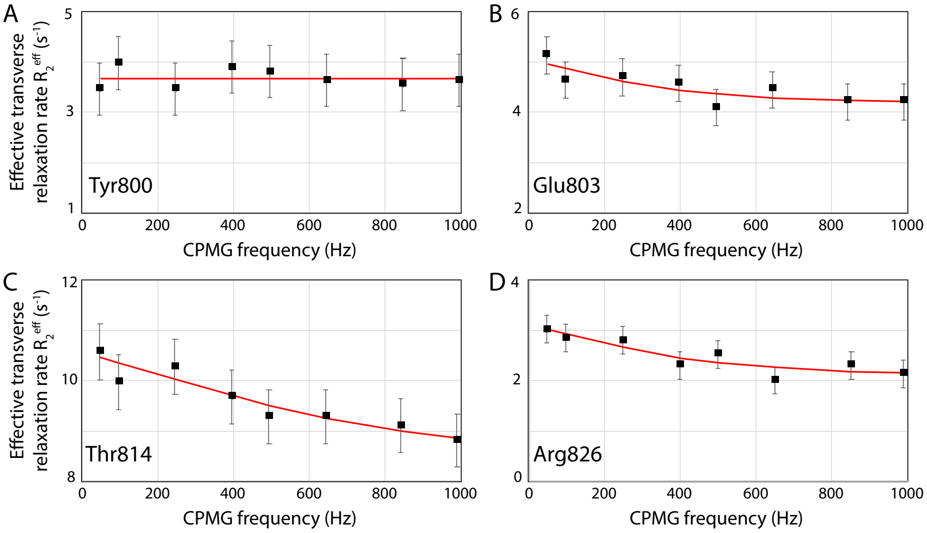

**Supplementary Figure S5. Microsecond-millisecond motions in BCL11A.** Carr-Purcell-Meiboom-Gill $R_2$ relaxation dispersion curves for residues Tyr800 (A), Glu803 (B), Thr814 (C) and Arg826 (D) of ZnF456 in the absence of DNA. Tyr800 is shown as a negative control. Glu803, Thr814 and Arg826 are the only residues where μs-ms are detected. Fit curves from the "LM63" model[1] are shown in red. Fitting equation is:

$$R_2{}^{eff} = R_2{}^0 + \frac{\Phi_{ex}}{k_{ex}}\left(1 - \frac{4\nu_{CPMG}}{k_{ex}}\tanh\left(\frac{k_{ex}}{4\nu_{CPMG}}\right)\right)$$

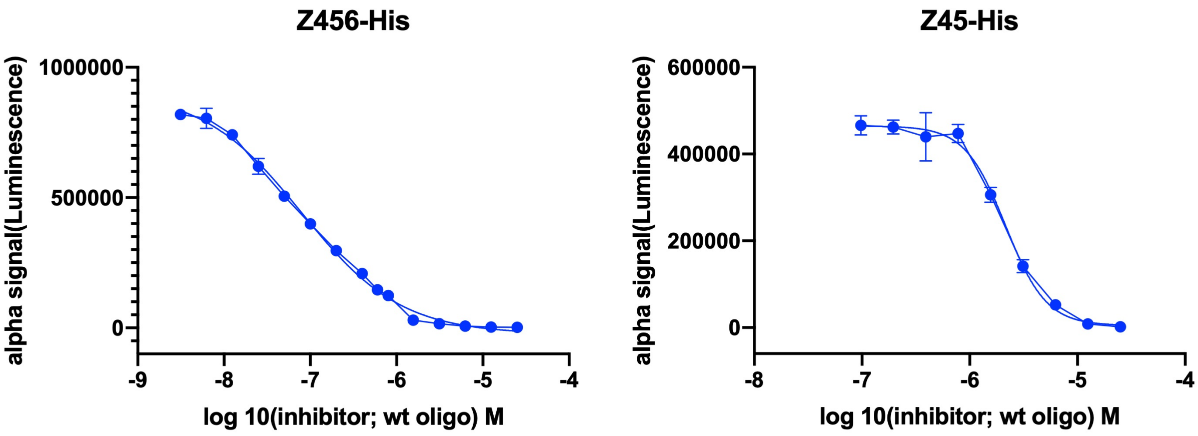

**Supplementary Figure S6. AlphaScreen assay for DNA binding affinity.** AlphaScreen determined $IC_{50}$ values of 79 ± 5 nM and 2.1 ± 0.2 μM for ZnF456 and ZnF45 binding to a 14-bp oligo, respectively.

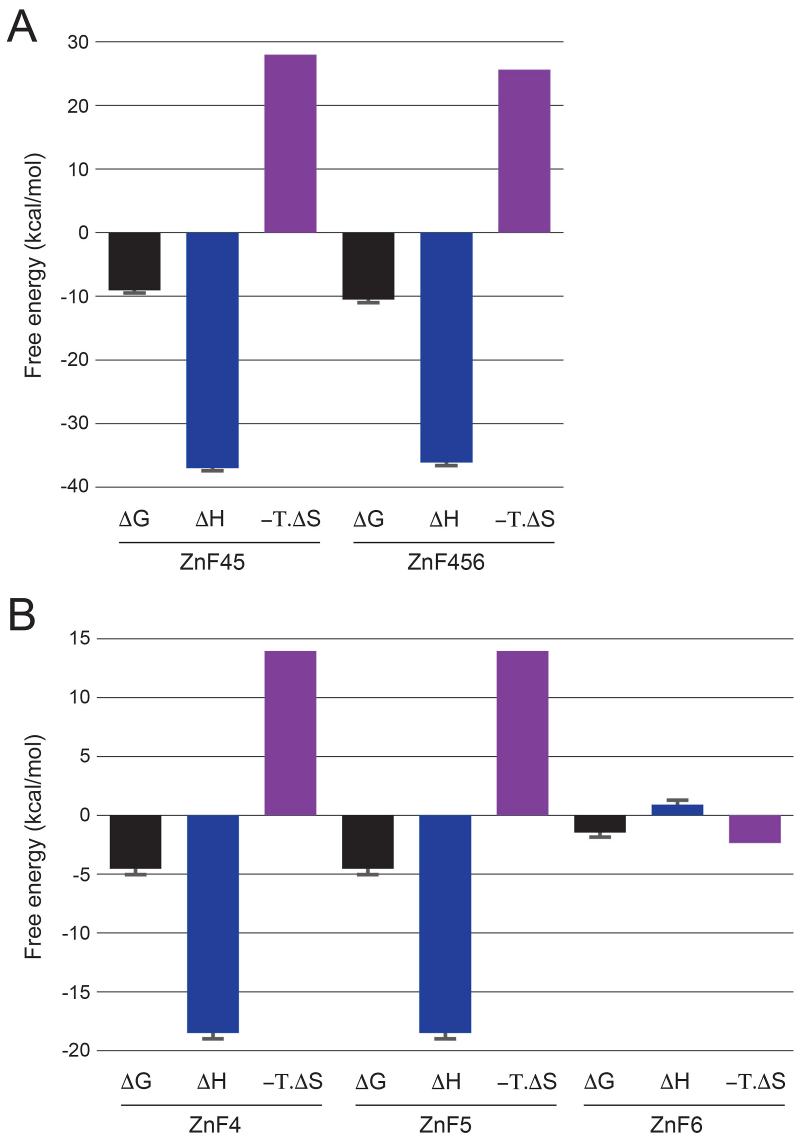

**Supplementary Figure S7. Summary of thermodynamic parameters extracted from ITC data.** A) Free energy, enthalpy and entropy extracted directly from ZnF45 and ZnF456 ITC data. B) Parameters calculated for each Zn finger assuming additivity and equal contributions of 4 and 5.

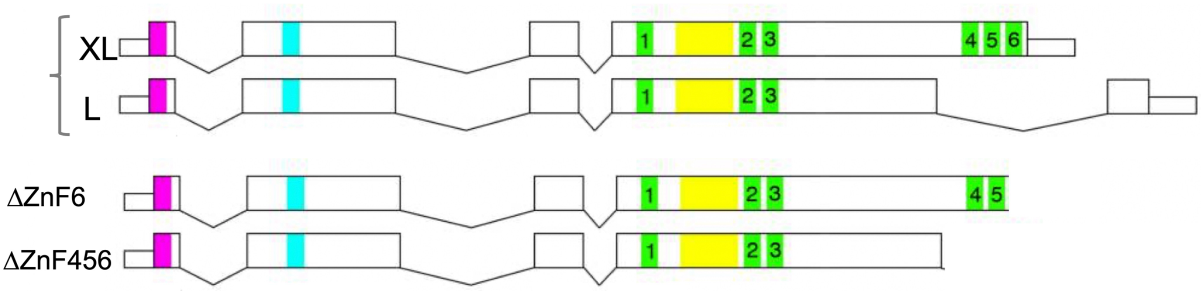

WT
CAGGCCCAGCTCAAAAGAGGGCAGACGCAG**CGACACTTGTGAGTACTGTGGGAAAGTCTTCAAGAACTGTAGCAATCTC**
**ACTGTCCACAGGAGAAGCCACACGGGCGAAAGGCCTTATAAATGCGAGCTGTGCAACTATGCCTGTGCCCAGAGTAGCA**
**AGCTCACCAGGCACATGAAAACGCATGGCCAGGTGGGGAAGGACGTTTACAAATGTGAAATTTGTAAGATGCCTTTTAG**
**CGTGTACAGTACCCTGGAGAAACACATGAAAAAATGGCACAGTGATCGAGTGTTGAATAATGATATAAAAACTGAATA**
**GAGGTATATTAAT**ACCCTCCCTCACTCCCACTTGA

456 deletion (Residues 742-835 deleted)
CAGGCCCAGCTCAAAAGAGGGCAGACGCAGACCCTCCCTCACTCCCACTTGA

WT
GGGGAAGGACGTTTACAAATGT**GAAATTTGTAAGATGCCTTTTAGCGTGTACAGTACCCTGGAGAAACACATGAAA**
**AAATGGCACAGTGATCGAGTGTTGAATAATGATATAAAAACTGAATAGAGGTATAT**TAATACCCTC

Znf6 del (Residues 803-835 deleted)
GGGGAAGGACGTTTACAAATGTTAATACCCTC

**Supplementary Figure S8. Structure of ZnF456 and ZnF6 BCL11A deletion alleles.** XL and L represent the major splice forms of BCL11A. The DNA sequences of the wild-type and deletion alleles are shown below.

**Supplementary movies**

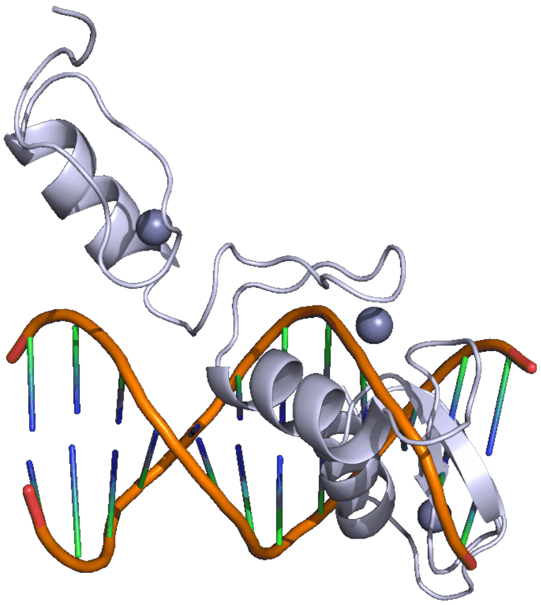

**Supplementary Movie S1. Morphing movie between the solution structure of BCL11A ZnF456** (ZnF6 would clash with DNA, light blue) **and its DNA-bound crystal structure** (wheat color).

**References**:

1. Luz, Z., and Meiboom, S. (1963). Nuclear Magnetic Resonance Study of the Protolysis of Trimethylammonium Ion in Aqueous Solution—Order of the Reaction with Respect to Solvent. The Journal of Chemical Physics *39*, 366-370. 10.1063/1.1734254.
